# Single-molecule long-read methylation profiling reveals regional DNA methylation regulated by Elongator Complex Subunit 2 in Arabidopsis roots experiencing spaceflight

**DOI:** 10.1101/2022.11.09.515199

**Authors:** Mingqi Zhou, Alberto Riva, Marie-Pierre L. Gauthier, Michael P. Kladde, Robert J. Ferl, Anna-Lisa Paul

## Abstract

**Background:** The Advanced Plant Experiment-04 - Epigenetic Expression (APEX04-EpEx) experiment onboard the International Space Station examined the spaceflight-altered cytosine methylation in two genetic lines of *Arabidopsis thaliana*, wild-type Col-0 and the mutant *elp2-5,* which is deficient in an epigenetic regulator Elongator Complex Subunit 2 (ELP2). Whole-genome bisulfite sequencing (WGBS) revealed distinct spaceflight associated methylation differences, presenting the need to explore specific space- altered methylation at single-molecule resolution to associate specific changes over large regions of spaceflight related genes. To date, tools of multiplexed targeted DNA methylation sequencing remain limited for plant genomes.

**Results:** To provide methylation data at single-molecule resolution, Flap-enabled next-generation capture (FENGC), a novel targeted multiplexed DNA capture and enrichment technique allowing cleavage at any specified sites, was applied to survey spaceflight-altered DNA methylation in genic regions of interest. The FENGC capture panel contained 108 targets ranging from 509 to 704 nt within the promoter or gene body regions of gene targets derived from spaceflight whole-genome data sets. In addition to genes with significant changes in expression and average methylation levels between spaceflight and ground control, targets with space-altered distributions of the proportion of methylated cytosines per molecule were identified. Moreover, trends of co-methylation of different cytosine contexts were exhibited in the same DNA molecules. We further identified significant DNA methylation changes in three previously biological process- unknown genes, and loss-of-function mutants of two of these genes (named as *EMO1* and *EMO2* for *ELP2-regulated Methylation in Orbit 1* and *2*) showed enhanced root growth rate.

**Conclusions:** FENGC simplifies and reduces the cost of multiplexed, targeted, single-molecule profiling of methylation in plants, providing additional resolution along each DNA molecule that is not seen in population-based short-read data such as WGBS. This case study has revealed spaceflight-altered regional modification of cytosine methylation occurring within single DNA molecules of cell subpopulations, which were not identified by WGBS. The single-molecule survey by FENGC can lead to identification of novel functional genes. The newly identified *EMO1* and *EMO2* are root growth regulators which may be epigenetically involved in plant adaptation to spaceflight.

## Background

DNA methylation can impact chromatin structure to epigenetically modulate gene expression in eukaryotes. In plants, DNA methylation plays a substantial role in the physiological adaptation to environmental changes and stresses [1]. Spaceflight presents a unique physiological challenge for plants, constituting an environment that is outside the evolutionary experience of any terrestrial organism. Plants will play a key role in the human life support systems envisioned for spaceflight and planetary exploration habitats [2,3], thus it is important to understand the nature of the responses plants mount in their physiological adaptation to this novel environment. Epigenetic change plays a crucial role in environmental adaptation, including adaptation to spaceflight [4, 5]. The Advanced Plant Experiment-04 - Epigenetic Expression (APEX- 04) experiment onboard the International Space Station (ISS) examined the spaceflight- altered cytosine methylation in *Arabidopsis thaliana* (Arabidopsis) using *elp2-5*, a mutant of an epigenetic regulator, Elongator Complex Subunit 2 (ELP2), along with Col- 0, a wild-type control [4]. RNA sequencing (RNA-seq) and Whole genome bisulfite sequencing (WGBS) revealed remodeled transcriptomes and DNA methylomes during spaceflight, remodeling that was regulated by ELP2. The spaceflight *elp2-5* phenotype was distinct from the space-grown wild-type plants in that root growth did not present a typical spaceflight directionality, which orients root growth away from the germinating seed, but rather displayed a distinct clumping of roots clustered around the seed. The molecular phenotype exhibited distinct differences in the transcriptome and methylome compare to the ground controls, with a majority of differentially expressed genes in *elp2- 5* roots being also differentially methylated between spaceflight samples and ground controls [4].

However, the short-read sequencing used in WGBS precludes the ability to phase regional epigenetic signatures along individual molecules such as consecutive methylated cytosines relative to transcription factor binding, nucleosome positioning, histone incorporation and modification [1, 6]. Furthermore, detection of differentially methylated cytosines (DmCs) by WGBS is affected by sequencing depth and false positives due to overestimation of 5mC levels [7], and many downstream analyses are restricted to DmCs [8, 9].

Flap-enabled next-generation capture (FENGC) is a novel method for sensitive, strand-specific capture of long DNA sequences from user-defined target regions for subsequent genotyping and detection of DNA modification [10]. In the single-tube FENGC protocol, standard oligonucleotides are designed to form 5’ flaps at any desired base pair position for cost-effective, specific cleavage of target sequences by flap endonuclease. In contrast to WGBS, targeted sequencing increases coverage depth, thereby improving data quality and further maximizing cost savings by limiting off-target sequencing. FENGC also reports 5mC over long, contiguous single molecules, and therefore readily identifies distinct epigenetic configurations of extended regions that cannot be distilled from short WGBS reads. Furthermore, because 5mC does not affect flap endonuclease activity and target sequence capture by FENGC occurs prior to deamination of C to U, target sequence primers are designed irrespective of DNA methylation [10, 11]. Hence, FENGC is not subject to the primer design constraints incumbent to bisulfite sequencing, namely, avoidance of potentially methylated sites or insertion of degenerate bases. Therefore, FENGC offers unprecedented latitude in targeted sequencing, especially for surveys of plant genomes that possess 5mC in three cytosine contexts, CG, CHG, and CHH [12]. The accuracy and efficiency of enzymatic methyl-seq, which is deployed in the methylation detection procedure of FENGC, has been verified in plants [7]; however, FENGC has yet to be applied to methylation analysis in plants.

Therefore, we set out to demonstrate the effective application of FENGC in plant methylation contexts and its power in identification of functional loci. The observed phenotypical and molecular alteration in roots of Col-0 and *elp2-5* grown on the ISS in APEX-04 present a short-read methylation dataset for a comparison with methylation profiles assayed by FENGC single-molecule long reads. WGBS-unidentified differential modification of DNA methylation substrates in functionally-annotated genes as well as unknown genes of plants grown in orbit have been detected, which led to identification of novel regulators of root growth.

## Results and Discussion

### Quality of FENGC assay

In the FENGC assay, strand cleavage was directed by precise targeting of flap endonuclease 1 (FEN1) to 5’ flaps formed by sequence-specific oligonucleotides, avoiding reliance on restriction sites or other specific motifs. The oligo panel targeted 108 regions in total, with 107 regions within the promoter or gene bodies of 86 genes of interest derived from our APEX-04 spaceflight and related experiments. These genes were differentially expressed in one or more spaceflight-associated environment(s) [4, 5, 13, 14]. A region of AT2G46830 with nearly no detectable 5mC in the APEX-04 WGBS experiment was included as an internal control to gauge efficient conversion and to rule out 5mC false positives. The captured set of DNA strands ranged from 509 to 704 nt (Additional file 1: Table S1), with mean of 580 nt and median of 572 nt. The sequence capture was performed as a single-tube multiplex reaction followed by enzymatic methyl-PCR (oligos listed in Additional file 2: Table S2). The FENGC products amplified from each reaction were subjected to PacBio long-read HiFi sequencing. For APEX-04 samples, including Ground control Col-0 Roots (GCR), Flight Col-0 Roots (FCR), Ground control *elp2-5* Roots (GER) and Flight *elp2-5* Roots (FER), the reads from the sequenced capture libraries were consistently ∼90% on-target (Figs. 1A and 1B; Additional file 3: Table S3). After filtering and combining the four biological replicates from each sample, on average, 82% of the 108 targeted regions were detected by at least 10 reads, with median coverage of 144, 612, 211 and 791x in four samples, respectively (Fig. 1C; Additional file 4: Table S4). The GC content of the 108 targets, ranging from 19% to 52%, was positively correlated with the aligned reads number (Fig. 1D; Additional file 5: Fig. S1), whereas there was a modest negative correlation in FENGC targets with an average GC content above 50% [10]. This indicates that both high and low GC content of targets influenced the reads yield of FENGC products.

**Fig. 1.**
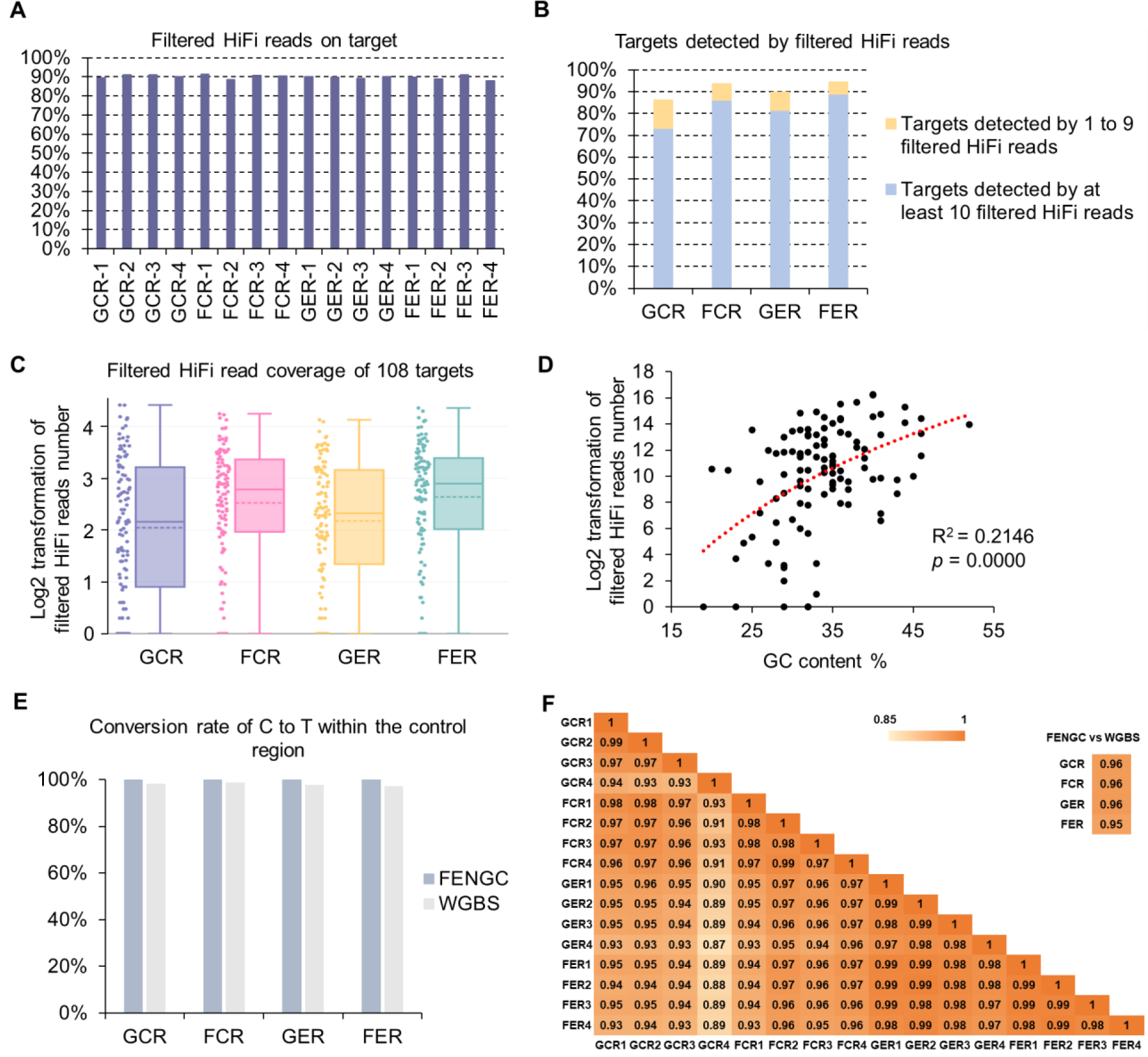
Application of flap-enabled next-generation capture (FENGC) in root samples of *Arabidopsis* experiencing spaceflight. (A) Percentage of filtered HiFi reads on target for all 108 targets. Reads with more than three consecutive methylated CHH sites were removed. GCR, Ground Col Roots; FCR, Flight Col Roots; GER, Ground control *elp2-5* Roots; FER, Flight *elp2-5* Roots. (B) Percentage of targets detected by at least 1 filtered HiFi read or at least 10 filtered HiFi reads. (C) Filtered HiFi read coverage of 108 targets. Log transformation of reads number was done by log2(read number +1). In boxplots, central solid line indicates median; central dot line indicates mean; box limits indicate upper and lower quartiles; whiskers indicate upper/lower adjacent values. Data dots are shown beside the boxplots. (D) Correlation between numbers of total filtered HiFi reads of all samples aligned to targets and the GC content of target regions. R-squared value and *p* value of logarithmic regression analysis are shown. (E) Conversion rates of C to T in sequencing results within the internal control region of 558 bp in AT2G46830 promoter. The region coordinates are in Additional file 4: Table S4. (F) Correlation coefficients between biological replicates of FENGC and between FENGC and WGBS for average cytosine methylation levels in seven example targets.

The efficiency of C to U conversion by EM-seq and hence quality of 5mC detection was calculated, using all cytosines from 558 bp of the internal control region of AT2G46830 promoter (Additional file 1: Table S1; Fig. 1E). The EM-seq conversion rates of the four samples pooled from the four biological replicates were all above 99%, on par with that of WGBS from 97% to 99% (Fig. 1E). In this study, we analyzed single molecules from seven target genes of interest, containing both hyper- and hypomethylated cytosines (described below). Collectively, levels of 5mC across CG, CHG, and CHH motifs exhibited correlation coefficients above 0.93 between any two biological replicates and above 0.95 between FENGC and WGBS results of APEX04 samples (Fig. 1F). The high reproducibility of FENGC targeted sequencing and strong agreement with WGBS results at diverse regions validate the FENGC application.

This case demonstrates multiple advantages for *in vitro* targeted measurement of 5mC in plant genomes by FENGC, including flexible target design, high sequencing depth, scalability, outstanding on-target ratio, and low cost (Table 1). To date, targeted DNA methylation detection in plants has predominantly relied on bisulfite sequencing of PCR amplicons. Primer design for bisulfite sequencing requires inclusion of degenerate bases to cover all possible methylation configurations of potential methylation sites that cannot be avoided, which can lead to assay bias and low multiplexing capacity.

**Table 1.**
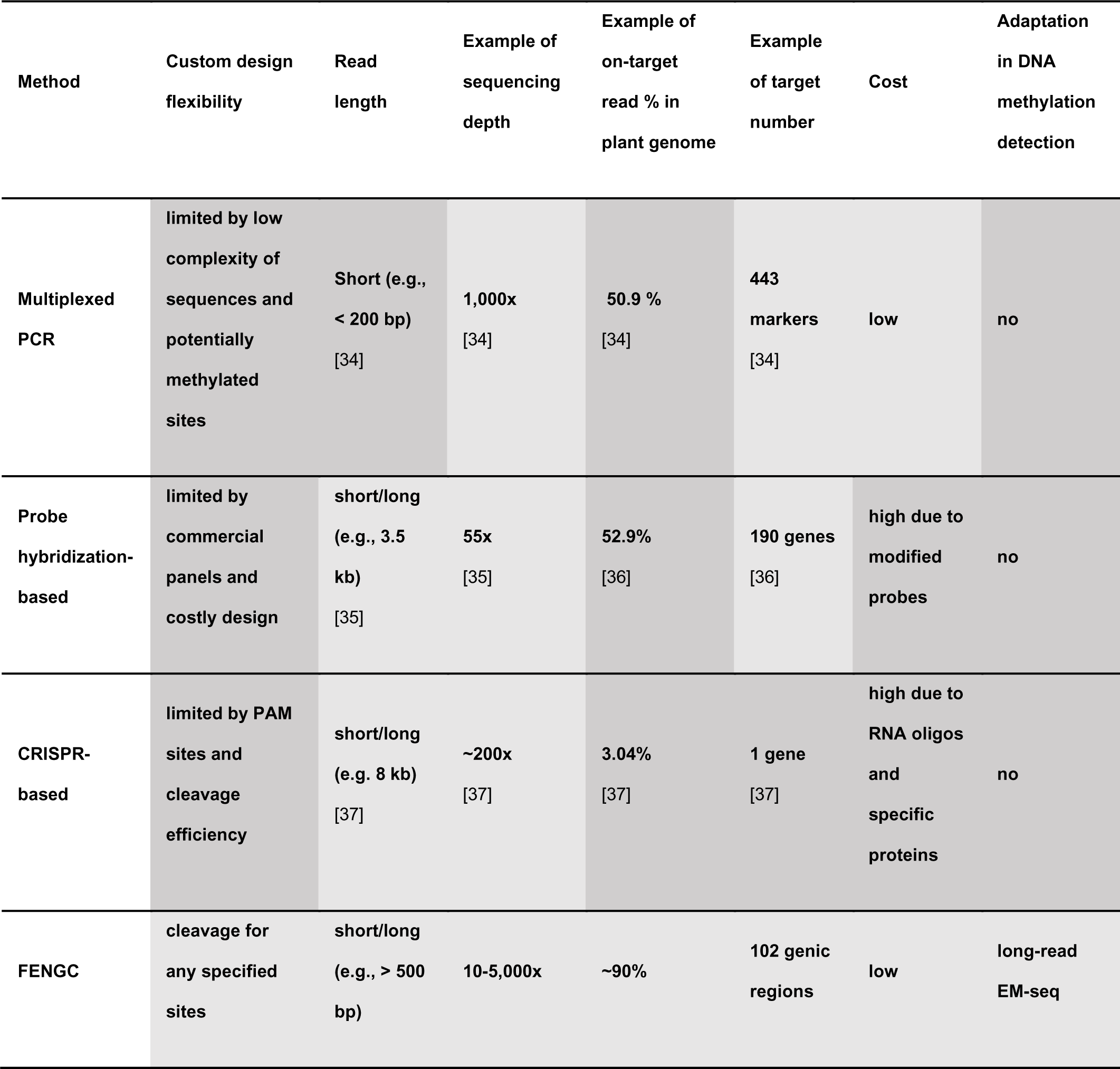
Comparison of multiplexed, targeted DNA sequencing methods in plants. Light gray indicates advantages and dark gray indicates disadvantages.

Recently, direct calling of DNA base modifications for plant sequences has extended the methods for profiling DNA methylation in plants [15, 16]; however, these applications require micrograms of input DNA and/or are limited to high-copy number sequences. The increasing need to examine large panels of specific single-copy loci, (as for differentially expressed genes of interest in an RNA-seq experiment) for epigenetic regulation by DNA methylation can pose substantial challenges, particularly when only nanograms of DNA are available and budgetary considerations are factors. FENGC overcomes these limitations, and therefore provides a unique approach that facilitates multiplexed, targeted DNA methylation analysis for plant research.

### Regional methylation detection by FENGC in comparison with WGBS data

One major advantage of long-read, single-molecule profiling of DNA methylation is assessment of 5mC at every cytosine along each contiguous DNA molecule, thereby presenting a coherent regional methylation status of an entire region from a single cell. In addition, the average 5mC levels across all molecules can be derived for comparison to WGBS data. For the present study, long-read, single-molecule sequencing identified gene targets showing not only average methylation level changes between spaceflight and ground control, but also space-altered distributions of the proportion of methylated cytosines per molecule.

Combining the four biological replicates, GCR, FCR, GER and FER samples yielded 79, 93, 88 and 96 target regions, respectively, with coverage of at least 10 reads (Additional file 3: Table S3 and Additional file 4: Table S4). The distributions of the proportion of methylated cytosines per molecule in each target region were calculated. Among them, 12 targets in 12 genes showed differential methylation proportion per molecule (DmPPM) for the comparison between spaceflight and ground control in Col-0 or *elp2-5* (Additional file 1: Table S1). Meanwhile, the APEX-04 WGBS-identified DmCs (methylation difference > 0.2, FDR < 0.01) between spaceflight and ground control were examined within 108 target regions of FENGC. Col-0 and *elp2-5* showed 8/2/7 and 5/2/2 targets possessing DmPPM for CG/CHG/CHH methylation, respectively. Among these regions, only one target with DmCHH for Col-0 and two targets with DmCHG for *elp2-5* were identified by WGBS (Additional file 1: Table S1). The comparison of detection in DmPPM and DmC demonstrated that the changes of contiguous methylation status observable in a specific subpopulation of molecules in FENGC may be indiscernible in a population-based analysis.

Next, we present four examples associated with defense responses to stimuli and three biological process-unknown genes regulated by ELP2 during plant growth in spaceflight to exhibit details of space-altered methylation status that are newly uncovered by FENGC (Additional file 1: Table S1). In each region, 93%-100% of the mapped reads are full-length, containing all CG/CHG/CHH sites of each target region (Additional file 5: Fig. S2). These results verify the high data quality for captured targets in the FENGC assay of space plants.

### Identification of previously unidentified space-altered DNA methylation in defense-related gene targets

Among 12 genes exhibiting DmPPM of spaceflight vs ground control, four defense-related genes showed space-triggered DNA methylation modification that was not discernible in WGBS. The first three examples were examined with CG methylation, for which DmPPM was shown in CG sites but no DmCG (methylation difference > 0.2, FDR < 0.01) between spaceflight and ground control was detected by WGBS in these target regions. *CML46* (AT5G39670), a gene encoding a calmodulin-like protein showing significant transcriptional down-regulation during spaceflight in Col-0 but not *elp2-5* plants, was assayed in a promoter region containing a MYB-like DNA binding element (Figs. 2A and 2B). Flight and ground control Col-0 roots (FCR and GCR), but not roots from *elp2-5* (FER and GER), showed a significant difference in the CG methylation level in an area flanking the MYB-like element, which contained five CG sites (Fig. 2C, in dashed frame). Consistent with these data, CG methylation proportion per molecule (mPPM) in this target region was significantly higher in FCR than GCR (Fig. 2D). A significant difference between GCR and GER was also observed. FENGC showed more molecules containing consecutively methylated CG sites in FCR (12%) compared to GCR (6.9%) in the area flanking this *cis*-regulatory element (Fig. 2E). Consistency between the levels of gene transcription, WGBS-derived population-based methylation status, and FENGC-derived mPPM, support the biological conclusion that ELP2 played a negative role in expression and a positive role in promoter methylation of *CML46* in spaceflight.

**Fig. 2.**
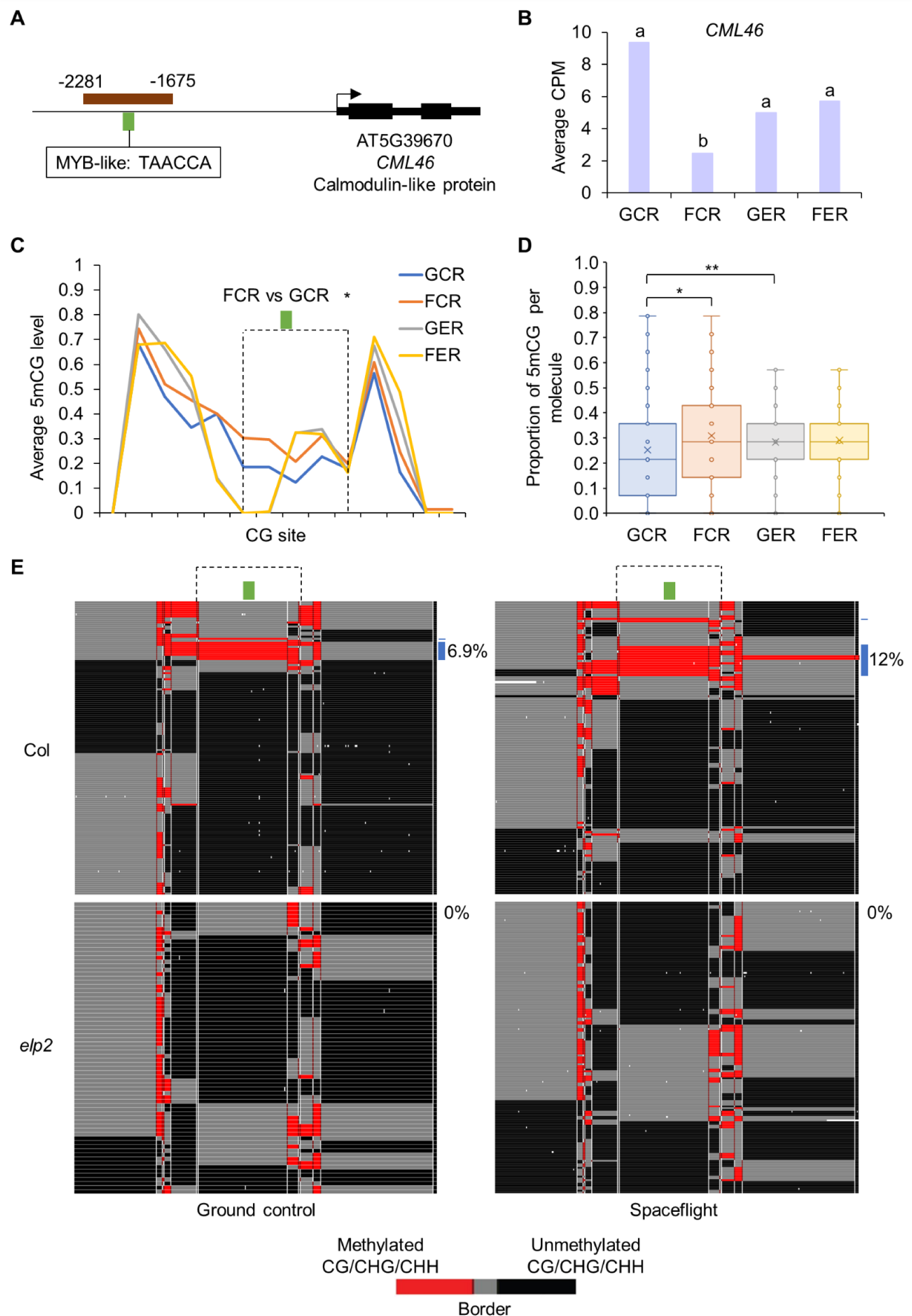
FENGC results for *CML46*. (A) Genic region of *CML46*. The target region is indicated by brown rectangle (target ID: AT5G39670-1 in S1 Table). A MYB-like binding element is indicated by green brick. The transcription start site is indicated by arrow. (B) Expression levels of *CML46* in each condition. Gene expression is indicated by average count per million mapped reads (CPM) in RNA-seq reported by Paul et al. 2021. Lowercase letters indicate significant differences between samples (FDR < 0.05 in DESeq2). (C) Average CG methylation level within the target region. The relative position of MYB-like binding element shown in (A) is indicated by green brick. The region indicated by dashed frame shows significant differences of CG methylation level between FCR and GCR (one-way ANOVA with post-hoc Tukey HSD test; *, *p* < 0.05). (D) Distribution of proportion of methylated CG sites per molecule in filtered, full-length HiFi reads aligned to the target region. The box indicates upper/lower quartiles and whisker indicates upper/lower adjacent values. The data points of mPPM are represented by dots. The line and “x” in the box indicate median and mean, respectively. The significant differences of methylation proportion are shown (Kruskal-Wallis test; *, *p* < 0.05; **, *p* < 0.01). (E) Heatmaps showing CG methylation of single molecules. Red indicates two or more consecutively methylated cytosines, black indicates two or more consecutively unmethylated cytosines, and gray indicates transitions between methylated and unmethylated cytosines. The reads from the four biological replicates (Additional file 4: Table S4) were combined to generate heatmaps for each sample. The relative position of MYB-like binding element shown in (A) and (C) is indicated by green brick. The region used for the CG methylation level comparison in (C) is indicated by dashed frame. Approximately 6.9% of reads in GCR and 12% of reads in FCR containing at least two consecutively methylated CG sites in both areas flanking the MYB-like binding element are labeled by blue bricks. One-way ANOVA with post-hoc Tukey HSD test was performed for pairwise comparisons of percentage of reads with this signature. FER vs FCR, *p* < 0.01.

*ABCB17* (AT3G28380), an ATP-binding cassette transporter gene that also showed a significantly lower transcription level in FCR than GCR, was assayed in a gene body region (Additional file 5: Figs. S3A and S3B). The overall patterns of averaged regional CG methylation levels were very similar between the four samples, which did not show significant changes between any pair of samples (Additional file 5: Fig. S3C); however, the CG mPPM in this target region was significantly higher in FCR than in GCR (Additional file 5: Fig. S3D). The FENGC single-molecule data showed that 3.7% of FCR reads possessed at least 6 consecutively methylated CG sites, whereas in GCR the percentage was 0.8% (Additional file 5: Fig. S3E). Biologically, *ABCB17* was repressed by spaceflight and ELP2 contributed to its transcription and gene body methylation.

Similarly, in the promoter region of *DEFL* (AT4G22217), a defensin-like gene (Additional file 5: Figs. S4A and S4B), the patterns of averaged CG methylation levels were highly similar and no significant alterations were detected between any two samples (Additional file 5: Fig. S4C); however, CG mPPM was significantly higher in FER than GER (Additional file 5: Figs. S4D and S4E). This higher methylation proportion correlates with the lower transcription level of this gene in FER compared to GER (Additional file 5: Fig. S4B). ELP2 enhanced expression and repressed promoter methylation of *DEFL* in spaceflight. These examples suggest that long-read FENGC single-molecule level methylation profiling can be more revealing of consequential methylation changes compared with population-based methylation methods such as WGBS, especially when alterations occur in a subpopulation of cells within a sample.

For the fourth example, a promoter fragment containing a MeJA-responsive element was selected in *PDF1.1* (AT1G75830), a gene coding a pathogenesis-related protein belonging to the plant defensin family and showing significant spaceflight induction in *elp2-5* (Figs. 3A and 3B). The average methylation level per site over all sequenced molecules showed significant differences at CHG and CHH sites between GER and GCR, as well as in CHH sites between FCR and GCR (Fig. 3C). For mPPM, significant differences were detected in all pairwise comparisons between flight conditions and genotypes at cytosines in all three methylation contexts (Fig. 3D). DmCs of WGBS between GER and GCR correlated with methylation changes detected by FENGC (Fig. 3D; Additional file 1: Table S1), while between spaceflight and ground control only three DmCGs in *elp2-5* were identified by WGBS, and the FENGC determined methylation level changes were smaller than 0.2 at these three sites (Fig. 3C), which was in line with the WGBS overestimation of methylation level compared with EM-seq, especially at CHG sites [7]. The patterns of methylation proportion in DNA molecules demonstrated that three cytosine contexts showed a trend of co-methylation in this region among cells in the population (Additional file 5: Fig. S5). Within each molecule, methylation proportion of CG is higher than CHG in 97.9% reads, and that of CHG is higher than CHH in 87.5% reads (Fig. 4A). Although three types of cytosine methylation exhibited a trend of coordination in the whole cell population, they could spread independently within subpopulations of cells. The CG methylation proportions were above 0.57 in all reads of GCR and above 0.28 in GER, while FCR and FER had reads with CG methylation proportion as low as 0. These data indicate that more molecules were hypomethylated in this region in *elp2-5* compared with Col-0 and in spaceflight compared with ground control.

**Fig. 3.**
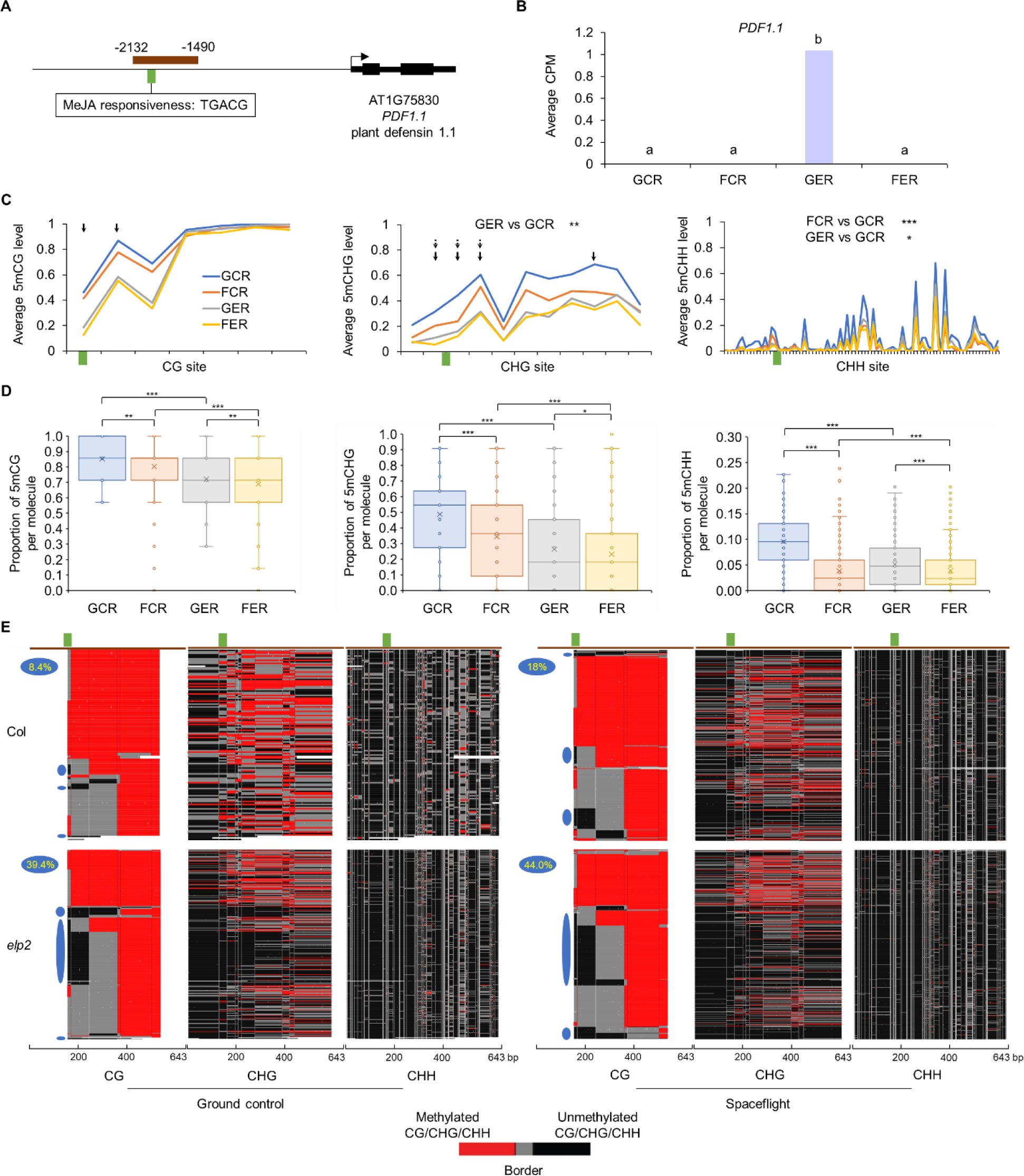
FENGC results for *PDF1.1*. (A) Genic region of *PDF1.1*. The target region is indicated by brown rectangle (target ID: AT1G75830-1 in Additional file 1: Table S1). A MeJA-responsive element is indicated by green brick. The transcription start site is indicated by the arrow. (B) Expression levels of *PDF1.1* in each condition. (C) Average CG, CHG or CHH methylation levels over all sequenced molecules of the target region. The relative position of MeJA-responsive element shown in (A) is indicated by green brick. The significant differences of methylation levels between samples for CHG and CHH sites in the target region are shown. One-way ANOVA with post-hoc Tukey HSD test; *, *p* < 0.05; **, *p* < 0.01; ***, *p* < 0.001. Black arrows indicate DmCs detected in WGBS using the threshold of methylation difference > 0.2 and FDR < 0.01. Arrows of solid line and dot line indicate DmCs of GER vs GCR and FER vs GER, respectively. (D) Distribution of proportion of methylated CG, CHG or CHH sites per molecule in filtered full-length HiFi reads aligned to the target region. The significant differences of methylation proportion are shown (Kruskal-Wallis test; *, *p* < 0.05; **, *p* < 0.01; ***, *p* < 0.001). (E) Heatmaps showing the cytosine methylation in single molecules. Reads from the four biological replicates (Additional file 4: Table S4) were combined to generate heatmaps for each sample. If the number of combined reads exceeded 1,000, 1,000 reads were randomly selected and plotted. The three heatmap panels show the patterns of methylation at CG, CHG, and CHH sites (vertical white lines) as indicated at bottom along the same DNA molecule in each row. The color legend is the same as that in Fig. 2E. Reads were hierarchically clustered using Kendall’s Tau based on CG methylation. The relative position of MeJA-responsive element shown in (A) is indicated by green brick. A footprint (blue ovals) with consecutive protection against methylation of the first 2 CG sites in the area flanking the MeJA-responsive element is present in 8.4%, 18.0%, 39.4% and 44.0% molecules in GCR, FCR, GER and FER, respectively. One-way ANOVA with post-hoc Tukey HSD test was performed for pairwise comparisons of percentage of reads with this footprint. FCR vs GCR, *p* < 0.05; GER vs GCR, *p* < 0.001; FER vs FCR, *p* < 0.001.

**Fig. 4.**
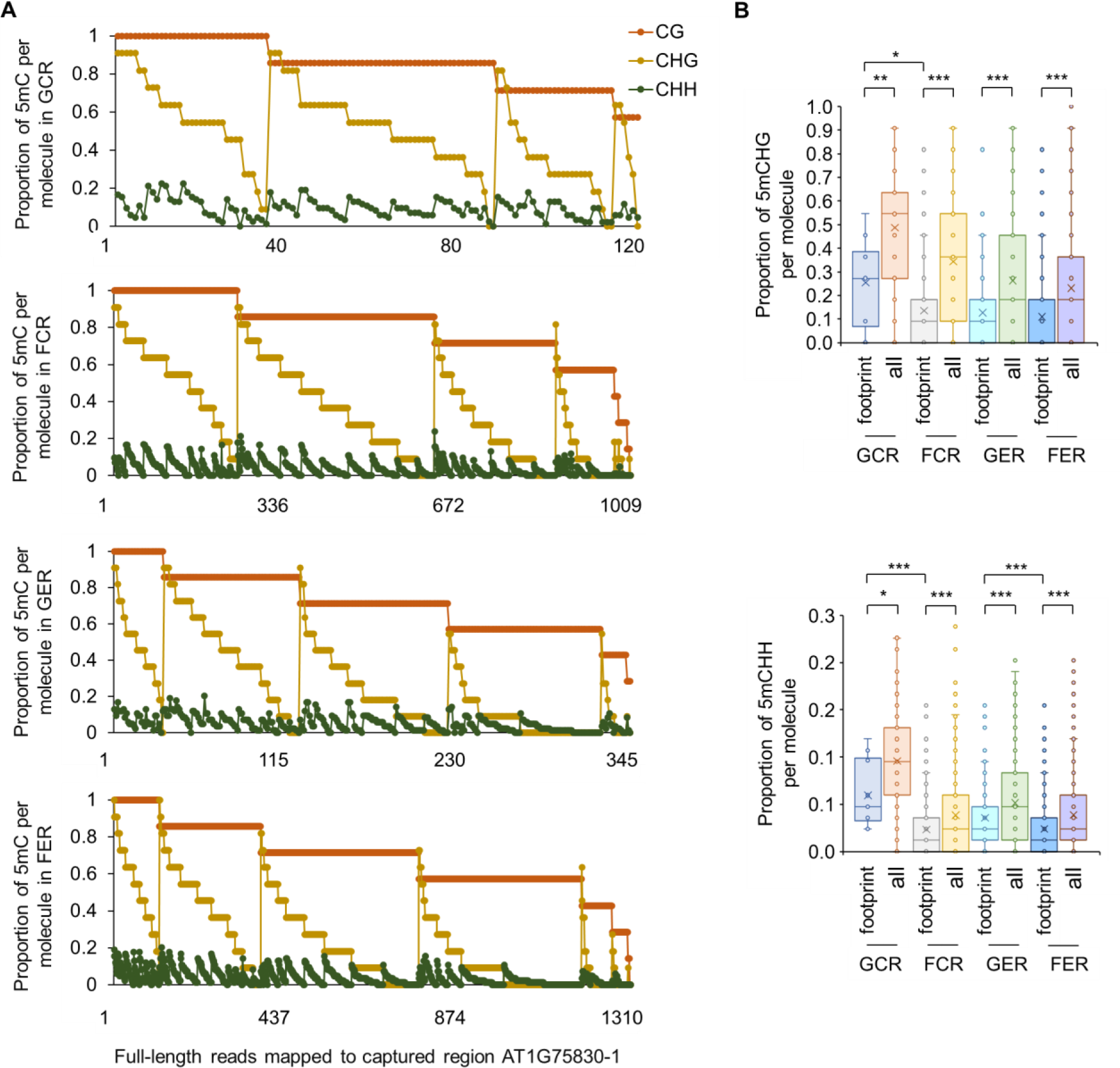
DNA methylation within same molecules in the target region of *PDF1.1*. (A) Proportion of methylated CG, CHG and CHH sites within the same molecule in the captured region of AT1G75830. There are 120, 1009, 345 and 1310 full-length reads covering all CG, CHG and CHH sites in this region (target ID: AT1G75830-1 in Additional file 1: Table S1), which are mapped for GCR, FCR, GER and FER samples, respectively. Ochre, gold and dark green dots indicate CG, CHG and CHH methylation proportions, respectively, in the same column, each representing one read. (B) Comparisons of methylation proportions per molecule in reads with a footprint and in all reads in the captured region of AT1G75830. CHG and CHH methylation in GCR, FCR, GER and FER samples are plotted, respectively. The footprint is defined as the first 2 CG sites in this target region that are both unmethylated. The significant differences of methylation proportion are shown (Kruskal-Wallis test; *, *p* < 0.05; **, *p* < 0.01; ***, *p* < 0.001).

The profiling of methylation data in this example also showcases another advantage of long-read, single-molecule FENGC assay of DNA methylation in plants, which is that all three methylation sequence contexts (CG/CHG/CHH) can be identified and plotted for the same molecule, thereby examining the entire methylation status of a gene region.

To determine the spatial distribution of these methylation differences, three separate but correlated panels were plotted for CG, CHG and CHH methylation in the same DNA molecule per row (Fig. 3E). In these plots, black areas are unmethylated regions, red areas are methylated regions, and gray areas are transition zones between methylated and unmethylated sites. It is visually apparent that spaceflight results in more unmethylated regions in Col-0, with the deepest loss of methylation concentrated near the MeJA-responsive element. It is also visually apparent that *elp2-5* shows a deeper loss of methylation in that region. For the CG methylation context, a footprint showing consecutive non-methylation of the first two CG sites in flanking area of the MeJA-responsive element was identified in 8.4%, 18.0%, 39.4% and 44.0% reads in GCR, FCR, GER and FER, respectively, quantifying the loss of CG methylation in this area. Interestingly, 60%, 86.5%, 97.5% and 97.6% reads with consecutive non-methylation in the second and the third CG sites in GCR, FCR, GER and FER, respectively, were also included in this footprint and showed consecutive non-methylation of the first three CG sites. In addition, CHG and CHH methylation proportions were significantly lower in the footprint area in the reads containing this footprint (Fig. 4B). These results are consistent with the previously observed high frequency of co-methylation/demethylation in nearby cytosines in Arabidopsis [17], and demonstrate that FENGC provides further opportunities for studying co-methylation of all three types of methylation along each single molecule.

Four targets shown here are associated with plant defense signaling, which is largely utilized in spaceflight adaptation of plants and the related genes can be involved in regulation of root growth phenotypes during spaceflight [18, 19]. It has also been reported that ELP2 modulates the epigenetic control of plant immune responses and defense processes [20]. The newly identified space-altered methylation in these targets is involved in ELP2-regulated defense/immunity-related pathways that may function in adaptation of plant roots grown in orbit.

### Identification of novel spaceflight-related epigenetic modification in biological process-unknown genes

The 12 genes with DmPPM between spaceflight and ground control were annotated using the Database for Annotation, Visualization and Integrated Discovery (DAVID, https://david.ncifcrf.gov/). Three genes were returned without gene ontology (GO) or functional annotation for biological process. Further investigation verified that the three target regions within these three genes possessed significant alteration between spaceflight and ground control in both average cytosine methylation level as well as mPPM in Col-0 or *elp2-5*. Moreover, loss-of-function mutants of two of these genes showed a root growth phenotype. We dubbed the two genes that could be epigenetically involved in plant adaptation to spaceflight under regulation of ELP2, AT1G27565 and AT5G52710, as *ELP2-regulated Methylation in Orbit* (*EMO*) *1* and *2*, respectively.

A promoter region was captured by FENGC for *EMO1* (AT1G27565), containing an anaerobic-responsive element (ARE) beside the transcription start site (TSS) (Fig. 5A). The presence of the ARE is consistent with the observed induction of hypoxia- responsive genes in spaceflight transcriptome data sets [5, 21]. *EMO1* was significantly up-regulated by spaceflight in *elp2-5*, but not in Col-0 (Fig. 5B). Regional average methylation level analysis detected significant differences between genotypes and between spaceflight and ground control in Col-0 only for CG sites (Fig. 5C). The calculation of mPPM showed significant alteration between spaceflight and ground control in all three types of cytosine context for Col-0 and in CG sites for *elp2-5* (Fig. 5D). Visualization of three types of methylated cytosines in each single molecule revealed footprints of methylation-free areas in areas in subpopulations of epialleles (Fig. 5E). Significantly, more molecules in *elp2-5* lacked cytosine methylation within this region than Col-0, and more molecules with methylation-free areas beside TSS were identified in spaceflight samples than ground control. These results were consistent with the observation of lower correlation of mPPM between GG and CHH in GCR than other samples (Additional file 5: Figs. S6 and S7). After demethylation initiated by spaceflight or mutation of ELP2 in a small proportion of cells, the co-methylation for different types of cytosine context in those molecules had been changed accordingly. Overall, ELP2 played a negative role in *EMO1* expression and maintained cytosine methylation of the *EMO1* promoter in a subpopulation of cells in roots. Spaceflight promoted demethylation in flanking areas of *EMO1* TSS, while the induction of *EMO1* transcription in space was repressed by ELP2.

**Fig. 5.**
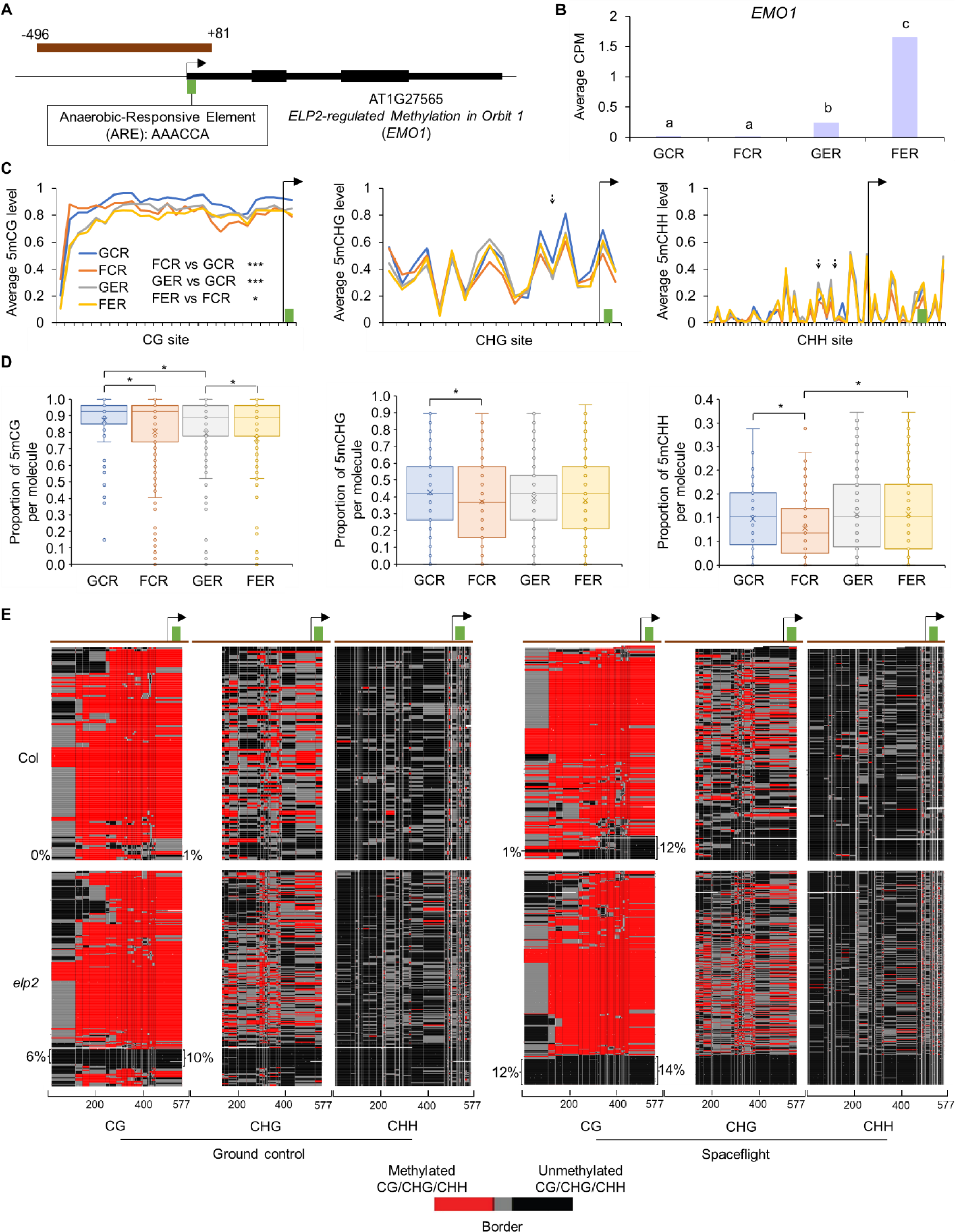
FENGC results for *EMO1*. (A) Genic region of *EMO1*. The target region is indicated by brown rectangle (target ID: AT1G27565-1 in Additional file 1: Table S1). An anaerobic-responsive element (ARE) is indicated by green brick. The transcription start site (TSS) is indicated by arrow. (B) Expression levels of *EMO1* in each condition. (C) Average CG, CHG or CHH methylation levels in cytosines within the target region. The relative position of ARE and TSS shown in (A) are indicated by green brick and arrow, respectively. The significant differences of CG methylation level between samples in the target region are shown, in which the first CG site is omitted. One-way ANOVA with post-hoc Tukey HSD test; *, *p* < 0.05; ***, *p* < 0.001. Black arrows indicate DmCs of FER vs GER detected in WGBS using the threshold of methylation difference > 0.2 and FDR < 0.01. (D) Distribution of proportion of methylated CG, CHG or CHH sites per molecule in filtered full-length HiFi reads aligned to the target region. The significant differences of methylation proportion are shown (Kruskal-Wallis test; *, *p* < 0.05). (E) Heatmaps showing the cytosine methylation in single molecules. The heatmap panels and color legend are the same as that in Fig. 3E. Reads were hierarchically clustered using Euclidean distance based on CG methylation. The percentages of reads without methylated CG sites and reads with at least 9 consecutive unmethylated CG sites at the 3’ end are shown on the left and right side of the CG panel, respectively. One-way ANOVA with post-hoc Tukey HSD test was performed for pairwise comparisons. For percentages of reads without methylated CG sites, FER vs GER, *p* < 0.05; GER vs GCR, *p* < 0.001; FER vs FCR, *p* < 0.01. For percentages of reads with at least 9 consecutive unmethylated CG sites at the 3’ end, FCR vs GCR, *p* < 0.01; GER vs GCR, *p* < 0.01.

The *EMO2* (AT5G52710) gene belongs to copper transport protein family [22]. The captured promoter region contains two MeJA-responsive elements (Fig. 6A). The transcription level of *EMO2* was lower in *elp2-5* than Col-0, and lower in spaceflight than ground control (Fig. 6B). Accordingly, the regional average methylation levels were significantly higher in *elp2-5* than Col-0 for CG and CHG sites, which was consistent with DmC analysis of WGBS (Fig. 6C). In contrast, CHH differential methylation levels detected by FENGC were smaller than that of WGBS. The mPPMs were higher in spaceflight than ground control in all three cytosine types for *elp2-5*, which explained the space-repressed expression of *EMO2* when ELP2 was deficient (Figs. 6B and 6D). Intriguingly, the areas of two MeJA-responsive elements both harbored a footprint of low CG methylation, whereas CHG sites localized within these elements showed methylation peaks, especially in *elp2-5* (Figs. 6C and 6E). When ELP2 was inactivated, the two CG methylation footprints were largely lost. Moreover, the correlations of mPPM between different types of cytosine context were lower in *elp2-5* than Col-0 (Additional file 5: Figs. S8 and S9). These results indicate the role of ELP2 in the maintenance of hypomethylated motifs and coordination of methylation for different cytosine types in this chromatin region, which could contribute to activation of *EMO2* transcription.

**Fig. 6.**
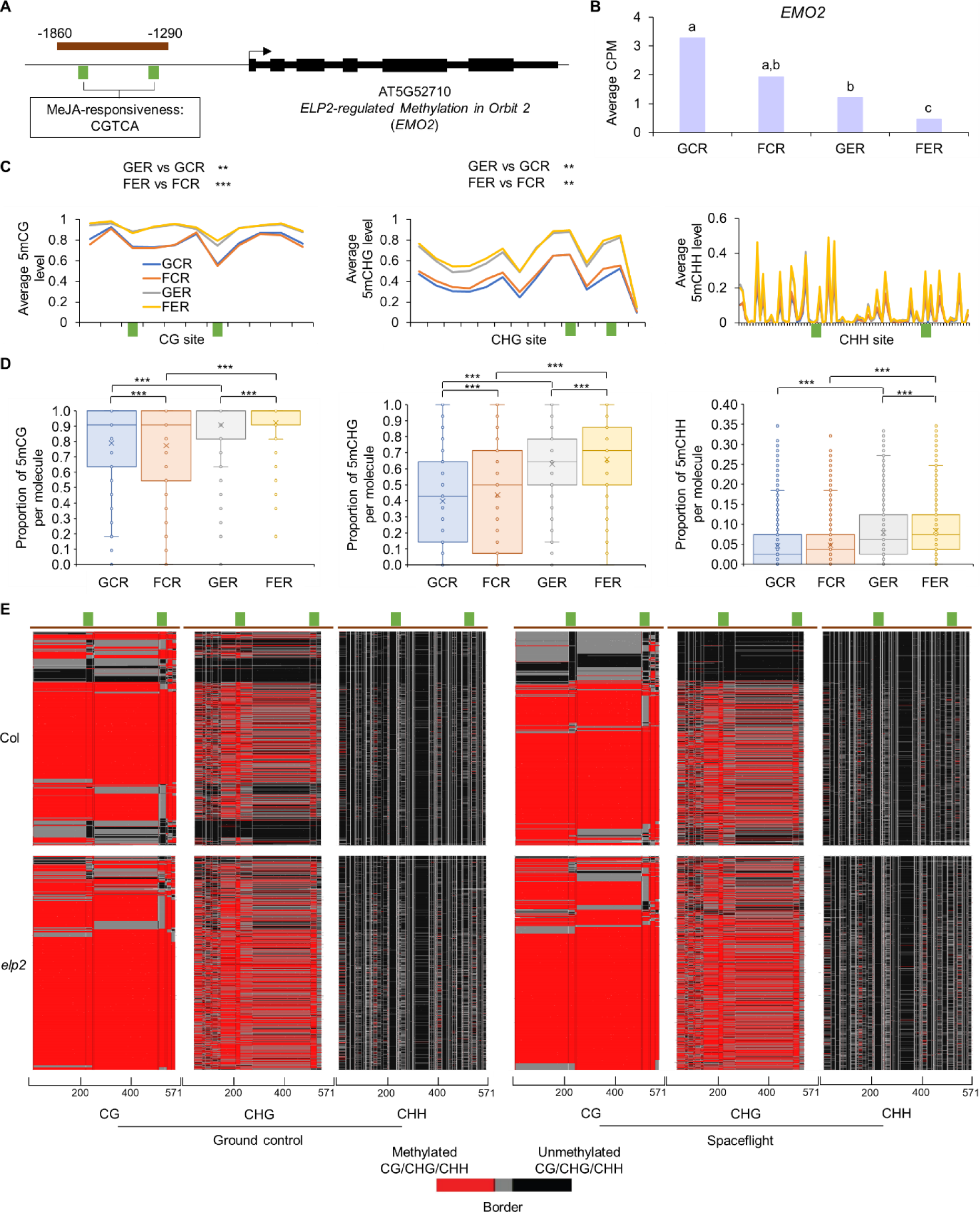
FENGC results for *EMO2*. (A) Genic region of *EMO2*. The target region is indicated by brown rectangle (target ID: AT5G52710 in Additional file 1: Table S1). Two MeJA-responsive elements are indicated by green bricks. The transcription start site (TSS) is indicated by arrow. (B) Expression levels of *EMO2* in each condition. (C) Average CG, CHG or CHH methylation levels in cytosines within the target region. The relative position of MeJA- responsive elements shown in (A) are indicated by green bricks. The significant differences of CG and CHG methylation levels between samples in the target region are shown. One-way ANOVA with post-hoc Tukey HSD test; **, *p* < 0.01; ***, *p* < 0.001. Black arrows indicate the DmCs detected in WGBS using the threshold of methylation difference > 0.2 and FDR < 0.01. Arrows with solid line, triangle, round end, and dot line indicate DmCs of GER vs GCR, FER vs FCR, FCR vs GCR, and FER vs GER, respectively. (D) Distribution of proportion of methylated CG, CHG or CHH sites per molecule in filtered full-length HiFi reads aligned to the target region. The significant differences of methylation proportion are shown (Kruskal-Wallis test; ***, *p* < 0.001). (E) Heatmaps showing the cytosine methylation in single molecules. The reads clustering, heatmap panels and color legend are the same as that in Fig. 5E.

The AT4G04990 gene encodes a serine/arginine repetitive matrix-like protein and the transcripts can be transported between roots and shoots [23]. The captured promoter region contains a TC-rich repeat that is related to defense and stress responses (Additional file 5: Fig. S10A). The transcription level of AT4G04990 was down-regulated by both spaceflight and mutation of ELP2 (Additional file 5: Fig. S10B). No DmC in WGBS or significant difference of average methylation level in FENGC was identified for this region, while DmPPMs of CG and CHH were detected between spaceflight and ground control in Col-0 (Additional file 5: Figs. S10C and S10D). In this region, a stronger correlation was observed between mPPMs of CHG and CHH than that of CG and CHH (Additional file 5: Figs. S10E and S11). Correlation between mPPMs of CG and CHG increased by spaceflight in Col-0 (Additional file 5: Fig. S11), which was partially due to space-induced demethylation of CG sites in a subpopulation of cells (Additional file 5: Figs. S10D, S10E and S12). The coordination of CG and non-CG methylation might be related to the transcription repression of AT4G04990 in spaceflight and *elp2-5*.

The features in alteration of DNA methylation at the single molecule level, such as DmPPM and methylation footprints, would be more informative for detection of epigenetic changes and better connect the regulation of gene expression to epigenetic modification, compared with population-based DmC levels that lose contextual epigenetic information of the single molecules. Therefore, FENGC data provide additional clues for further selection of target genes of interest for downstream analysis, especially when candidates include unknown genes.

### *EMO1* and *EMO2* are negative regulators of root growth

T-DNA insertion lines of the three biological process-unknown genes identified by FENGC were characterized. Two lines of each gene tested showed disrupted gene expression levels compared with Col-0 (Additional file 5: Fig. S13). Thus, SALK_092504 and SAIL_344_F07 were named as *emo1-1* and *emo1-2*, while SALK_140748C and SALK_091636C were named as *emo2-1* and *emo2-2*, respectively. In a constant LED light condition that was similar with Veggie facility onboard ISS, *emo1* and *emo2* mutants showed significantly enhanced root growth rates compared with Col-0 (Figs. 7A-7C). The primary root length of *emo1-1*, *emo1-2*, *emo2-1*, and *emo2-2* were significantly longer than that of Col-0 in 4-day and 7-day old seedlings. The two mutants of AT4G04990 exhibited similar growth phenotype with wild type. Meanwhile, treatments using 2-D clinorotation at 4 rpm or reorientation to 90° did not show altered gravitropic responses in roots of any line tested, although roots of *EMO1* and *EMO2* mutants still grew faster than Col-0 on the clinostat (Additional file 5: Figs. S14 and S15). These results demonstrated that *EMO1* and *EMO2* are two negative regulators of root growth, but not regulators of root gravitropism.

**Fig. 7.**
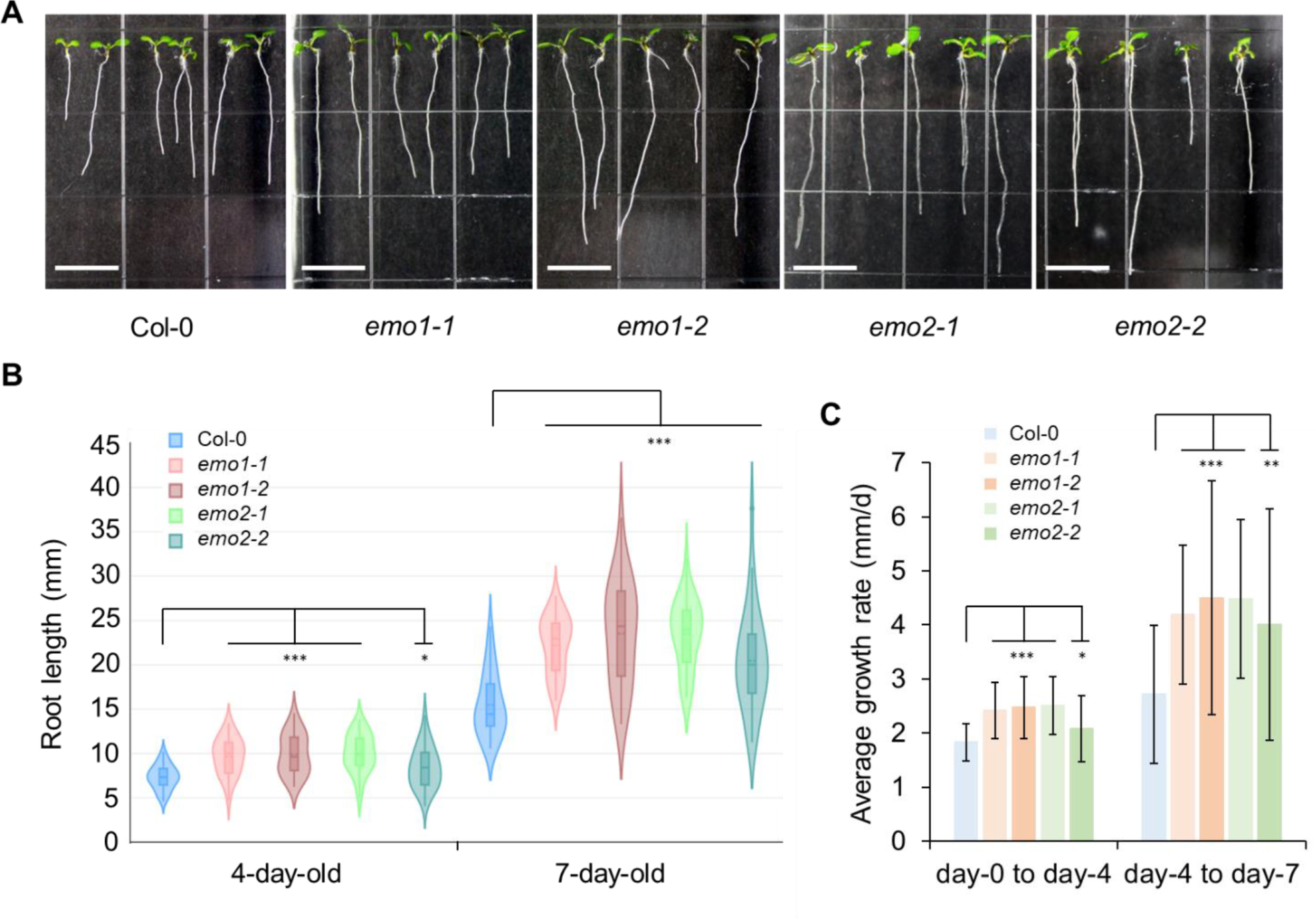
Enhanced root growth in mutants of *EMO1* or *EMO2*. (A) 7-day old seedlings of Col-0, *emo1-1*, *emo1-2*, *emo2-1*, and *emo2-2*. Scale bars indicate 10 mm. (B) Primary root length of 4-day old and 7-day old seedlings for each line. In violin/boxplots, central solid line indicates median; central dot line indicates mean; box limits indicate upper and lower quartiles; whiskers indicate upper/lower adjacent values. Kruskal-Wallis test; *, *p* < 0.05; ***, *p* < 0.001. (C) Average root growth rate (mm per day) from day-0 to day-4, and from day-4 to day-7. Student’s t test, *, *p* < 0.05; **, *p* < 0.01; ***, *p* < 0.001. Data ± SD, n = 55, 38, 33, 36, 34, 41, 38, 33, 36, 34 from left to right.

Remodeled plant root growth has been reported in orbital spaceflight [24]. Both direction of root tip growth and root growth rate were impacted in Arabidopsis plants grown on the ISS [18, 24, 25]. The primary root length of 8.5-day old seedlings for both Col-0 and Ws were significantly shorter in flight samples than ground control. This influence on root growth from spaceflight involves epigenetic and transcriptional modulation [4, 13]. As a epigenetic modulator of root growth and development, ELP2 modifies DNA methylation or histone acetylation of a range of transcription factors and other developmental regulators in roots [26]. On the ISS, the *elp2-5* mutant exhibited severe growth retardation in roots compared with Col-0 [4]. Here, *EMO1* and *EMO2* have been identified as two downstream targets of DNA methylation modification led by ELP2 and/or caused by extraterritorial environment. Together with the root growth rate phenotype in their mutants, *EMO1* and *EMO2* can play a role in root growth regulation during space adaptation of plants, which may involve the epigenetic signaling pathways controlled by ELP2. Moreover, although the mutants of AT4G04990 did not show root growth alteration on the ground, further investigation will be needed to examine its function in space.

## Conclusion

The targeted long-read, single-molecule profiling enabled by FENGC reveals detailed epigenetic modifications occurring in cell subpopulations and identifies epialleles present at low frequencies, whereas short-read WGBS only detects alteration of the average methylation level in cytosines. On one hand, FENGC and WGBS show consistency in the detection of cytosine methylation levels. On the other hand, FENGC enables single-molecule determination of methylation alteration in genes of interest defined by users, which greatly extends interpretation beyond that of WGBS. Analysis of DmPPM serves as critical addition to DmC data in the methylation survey. The capacity of target cleavage at any specified sites enables wide application of FENGC, such as studies on plant development, environmental responses, and generation of biomarkers. In the current work, FENGC enabled a richer contextual survey of regional methylation in genes of interest identified from primary transcriptomic and WGBS screening, successfully identifying two novel root growth regulators through analysis of DNA methylation alteration. In summary, FENGC provides the plant and space biology communities with a powerful tool for the practical, routine application of multiplexed targeted sequencing and coordinated methylation analysis at single-molecule resolution. This FENGC screen of spaceflight-associated genes revealed candidate genes with regulatory regions that harbored significant alterations in consecutive spans of methylation. Further genetic analysis identified *EMO1* and *EMO2*, which could be epigenetically modulated by ELP2 for both transcription and DNA methylation during spaceflight. The DmPPM of *EMO1* and *EMO2* between *elp2-5* and Col-0 indicated that ELP2-mediated cytosine modification in these two genic regions occurred in a specific subpopulation of cells in space. These two genes are novel candidates for further investigation of root growth regulatory signaling elicited by ELP2 in plant adaptation to spaceflight.

## Methods

### Plant materials and operations of spaceflight and ground controls

The *Arabidopsis thaliana* lines Col-0 and *elp2-5*, as well as spaceflight experimental operations, have been previously documented [4, 20]. The Advanced Plant Experiment 04 – Epigenetic Expression (APEX-04-EPEX) spaceflight experiment was launched onto International Space Station (ISS) through the SpaceX mission CRS-10. In brief, seeds were sterilized and sown on growth medium containing 0.5% phytagel, 0.5× Murashige–Skoog salts, 0.5% (w/v) sucrose, and 1 × Gamborg’s Vitamin Mixture, in Petri dishes (100 mm × 15 mm; Fisher Scientific, Pittsburgh, USA), which were then sealed with breathable tape (3M Micropore, Maplewood, USA). Plants were grown in the Vegetable Production System (Veggie) facility on the ISS and the Veggie hardware in the ISS environmental simulator (ISSES) chamber at Kennedy Space Center under constant LED light of 100–135 μmol/m^2^s Photosynthetically Active Radiation (PAR) until 11 days after germination, prior to harvest using RNAlater (Ambion, Grand Island, USA) in Kennedy Space Center fixation tubes (KFTs). Plants from each plate were collected into individual KFTs and stored at -80°C until delivery to the lab for analysis.

### Genomic DNA Isolation and flap-enabled next-generation capture (FENGC) assay

DNA extraction was conducted using the protocol designed for RNAlater preserved plant samples [27]. The same set of DNA samples from Col-0 and *elp2-5* root tissues used for whole-genome bisulfite sequencing (WGBS) [4] were subjected to flap-enabled next-generation capture (FENGC) assay. The samples included four biological replicates of each condition: Ground control Col-0 Roots (GCR), Flight Col-0 Roots (FCR), Ground control *elp2-5* Roots (GER) and Flight *elp2-5* Roots (FER).

FENGC oligo-1, -2 and -3 were designed using FENGC Oligonucleotide Designer (FOLD) program (https://github.com/uf-icbr-bioinformatics/FOLD), to bind target DNA to form 5′ flap structures that were cleaved by flap endonuclease (FEN), which enabled ligation between target fragments and universal adaptors [10]. There were 108 targets selected within promoter and gene body regions of 87 genes of interest derived from previous spaceflight-associated transcriptome and DNA methylome studies [4, 5, 13, 14]. The detailed information of the 108 targets is in Additional file 1: Table S1. All oligos used in this study are listed in Additional file 2: Table S2. All captured sequences are from the coding strand of each gene.

FENGC experiments were performed as described by Zhou et al. [10], with modifications to adapt to plant genome. In brief, approximately 200 ng of Arabidopsis genomic DNA was fragmented by sonication using a Bioruptor Pico (Diagenode, Denville, USA) for 6 cycles of 5 sec on and 30 sec off in 20 µl ddH2O. The concentration of oligo-1, -2, and -3 was 18 nM each in the master tube as recommended [10]. Flap cleavage was performed in the presence of FENGC universal oligo 1-T (U1-T oligo), flap oligos-1 and -2 specific for 108 targets, and thermostable FEN1 (New England Biolabs, Ipswich, USA). Ligation was initiated by adding a stock mixture of oligos-3 for all 108 targets, U2 oligo, and Ampligase (Lucigen, Middleton, USA) to the same tube.

DNA not ligated to U2 oligo and therefore not protected by 3’ C3 Spacer was removed by addition of exonucleases I and III (NEB, Ipswich, USA). The enriched products were purified and then subjected to enzymatic methylation conversion according to the manual of NEBNext Enzymatic Methyl-seq Conversion Module (NEB, Ipswich, USA).

Converted DNA was amplified for 23 cycles using U1 and U2 primers and KAPA HiFi HotStart Uracil+ DNA polymerase (Roche, Pleasanton, USA), according to the manufacturer’s instructions. FENGC amplification products were purified and product quality and quantity were determined using Agilent TapeStation D1000 system (Agilent, Santa Clara, USA) and Qubit 3 Fluorometer (Thermo Fisher, Waltham, USA), respectively.

### Pacific Biosciences Sequel Ile SMRT sequencing and data analysis

The library construction and sequencing of FENGC products on the Pacific Biosciences (PacBio) Sequel IIe SMRT platform was performed by Interdisciplinary Center for Biotechnology Research (ICBR) at the University of Florida. The HiFi reads were generated using 30-h movie of circular consensus (CCS) mode with filtering for ≥3 passes per single polymerase read, with default settings for other parameters. HiFi reads mapping and visualization of DNA methylation states of single molecules were performed using MethylMapper as previously described [28], with modifications to accommodate plant genomes. Reads with more than three consecutive methylated CHH sites were removed [29].

Demultiplexed CCS reads data files of PacBio Sequel IIe SMRT sequencing have been deposited with the accession number of PRJNA857037 to BioProject database (https://www.ncbi.nlm.nih.gov/bioproject/) in National Center for Biotechnology Information (NCBI), and with the OSD identifier of OSD-625 (DOI: 10.26030/rp5n-0d70) in NASA GeneLab.

### Roots characterization assay

The *Arabidopsis thaliana* T-DNA insertion lines SALK_092504 (*emo1-1*), SAIL_344_F07 (*emo1-2*), SALK_140748C (*emo2-1*), SALK_091636C (*emo2-2*), SAIL_205_A08, and WiscDsLox441G3 were obtained from Arabidopsis Biological Resource Center (https://abrc.osu.edu/). Seeds were sterilized, incubated in ddH2O at 4°C for 3 days, and then sown on 0.5x Murashige–Skoog medium in Petri dishes as described above. Plates were placed on vertical racks at 21°C in 24 h LED light condition at approximately 100 μmol/m^2^s PAR. The 4-day old were photographed. Then plants were subjected to three treatments. The first set of plates were kept quiescent and seedlings were grown to 7-day old and photographed. The second set of plates were mounted to a 2-dimentional clinostat to be rotated at 4 rounds per min until plants were 7-day old, and then photographed. The third set of plates were reoriented 90° to the horizontal, and photographed at 0 h, 6 h and 24 h after reorientation. Roots were traced and measured using JFilament plugin of ImageJ [30, 31]. Root growth measurements were processed using the R script RootMeasurement (https://github.com/eschultzphd/RootMeasurement/tree/master) [32]. The quantitative real-time PCR (qPCR) assay to verify gene knock-out or knock-down was performed using roots of 7-day old seedlings as previously described [33]. UBQ11 (AT4G05050) was used as the internal control and primers are listed in Additional file 2: Table S2.

### Availability of data and materials

Demultiplexed CCS reads data files of PacBio Sequel IIe SMRT sequencing have been deposited with the accession number of PRJNA857037 to BioProject database (https://www.ncbi.nlm.nih.gov/bioproject/) in National Center for Biotechnology Information (NCBI), and with the OSD identifier of OSD-625 (DOI: 10.26030/rp5n-0d70) in NASA GeneLab. Requests for updated versions of FENGC protocol, FOLD and MethylMapper scripts should be directed to Michael Kladde.

## Abbreviations

*ABCB17*: ATP-binding cassette B17
*ANOVA*: Analysis of Variance
*APEX-04-EPEX*: Advanced Plant Experiment 04 – Epigenetic Expression
*ARE*: Anaerobic-responsive element
*CCS*: Circular consensus sequencing
*CML46*: Calmodulin like 46
*CPM*: Count per million mapped reads
*Col-0*: Columbia-0
*DEFL*: Defensin-like
*DmC*: Differentially methylated cytosine
*DmPPM*: Differential methylation proportion per molecule
*ELP2*: Elongator complex subunit 2
*EMO*: ELP2-regulated methylation in orbit
*EM-seq*: Enzymatic methylation sequencing
*FCR*: Flight Col-0 roots
*FDR*: False discovery rate
*FEN1*: Flap endonuclease 1
*FENGC*: Flap-enabled next-generation capture
*FER*: Flight elp2-5 roots
*FOLD*: FENGC oligonucleotide designer
*GCR*: Ground control Col-0 roots
*GER*: Ground control elp2-5 roots
*GO*: Gene ontology
*HSD*: Honestly significant difference
*ICBR*: Interdisciplinary Center for Biotechnology Research
*ISS*: International Space Station
*ISSES*: ISS environmental simulator
*KFT*: Kennedy Space Center fixation tube
*Methyl-seq*: Methylation sequencing
*MYB*: Myeloblastosis
*mPPM*: Methylation proportion per molecule
*PacBio*: Pacific Biosciences
*PAR*: Photosynthetically Active Radiation
*PDF1.1*: Plant defensin 1.1
*qPCR*: Quantitative real-time polymerase chain reaction
*RNA-seq*: RNA sequencing
*TSS*: Transcriptional start site
*T-DNA*: Transfer DNA
*UBQ11*: Ubiquitin 11
*Veggie*: Vegetable Production System
*WGBS*: Whole-genome bisulfite sequencing
*5mC*: 5-Methylcytosine

## Acknowledgments

We thank members of UF Space Plants Lab for helpful discussion and assistance in experiments, as well as members of UF ICBR, NextGen DNA sequencing (RRID:SCR_019152), Bioinformatics (RRID:SCR_019120) and Gene Expression and Genotyping (RRID:SCR_019145) cores, for their support and services. We also thank Jason O. Brant for his assistance in developing the FOLD program.

## Author contributions

MZ, RJF and A-LP designed this study. MZ conducted the experiments and analyzed the data. MZ and MPK developed the FENGC methodology. AR, M-PLG and MPK contributed to the FENGC oligo design program. MZ, MPK, RJF and A-LP wrote the manuscript.

## Ethics approval and consent to participate

Not applicable.

## Consent for publication

Not applicable.

## Supplementary Information

Additional file 1: Table S1. Information of 108 capture targets.

Additional file 2: Table S2. Oligos used in this study.

Additional file 3: Table S3. Filtered HiFi reads on target and targets detected.

Additional file 4: Table S4. Number of filtered HiFi reads aligned to 108 targets.

Additional file 5: Figs. S1-S15.

**Fig. S1.** Correlation between numbers of filtered HiFi reads aligned to targets and the GC content of target regions in each sample. **Fig. S2.** Percentage of full-length reads and truncated reads mapped in seven selected target regions. **Fig. S3.** FENGC results for *ABCB17*. **Fig. S4.** FENGC results for AT4G22217. **Fig. S5.** Correlation between proportions of methylated cytosines in different sequence contexts within the same molecule in the captured region of *PDF1.1*. **Fig. S6.** Correlation between proportions of methylated cytosines in different sequence contexts within the same molecule in the captured region of *EMO1* (AT1G27565). **Fig. S7.** Proportion of methylated CG, CHG and CHH sites within the same molecule in the captured region of *EMO1* (AT1G27565). **Fig. S8.** Correlation between proportions of methylated cytosines in different sequence contexts within the same molecule in the captured region of *EMO2* (AT5G52710). **Fig. S9.** Proportion of methylated CG, CHG and CHH sites within the same molecule in the captured region of *EMO2* (AT5G52710). **Fig. S10.** FENGC results for AT4G04990. **Fig. S11.** Correlation between proportions of methylated cytosines in different sequence contexts within the same molecule in the captured region of AT4G04990. **Fig. S12.**

Proportion of methylated CG, CHG and CHH sites within the same molecule in the captured region of AT4G04990. **Fig. S13.** Relative transcription levels of *EMO1*, *EMO2* or AT4G04990 in their mutants, respectively. **Fig. S14.** Characterization of root growth in mutants of *EMO1*, *EMO2* or AT4G04990. **Fig. S15.** Root gravitropism in mutants of *EMO1*, *EMO2* or AT4G04990.

