## Additional file 5: Figs. S1-S15. for "Single-molecule long-read methylation profiling reveals regional DNA methylation regulated by Elongator Complex Subunit 2 in Arabidopsis roots experiencing spaceflight"

**Fig. S1.** Correlation between numbers of filtered HiFi reads aligned to targets and the GC content of target regions in four biological replicates of Ground control Col-0 Roots (GCR), Flight Col-0 Roots (FCR), Ground control *elp2-5* Roots (GER) and Flight *elp2-5* Roots (FER). Log transformation of reads number was done by log_2_(read number +1). R-squared value and *p* value for logarithmic regression analysis are shown. The aligned reads number was positively correlated to the GC content of target regions.

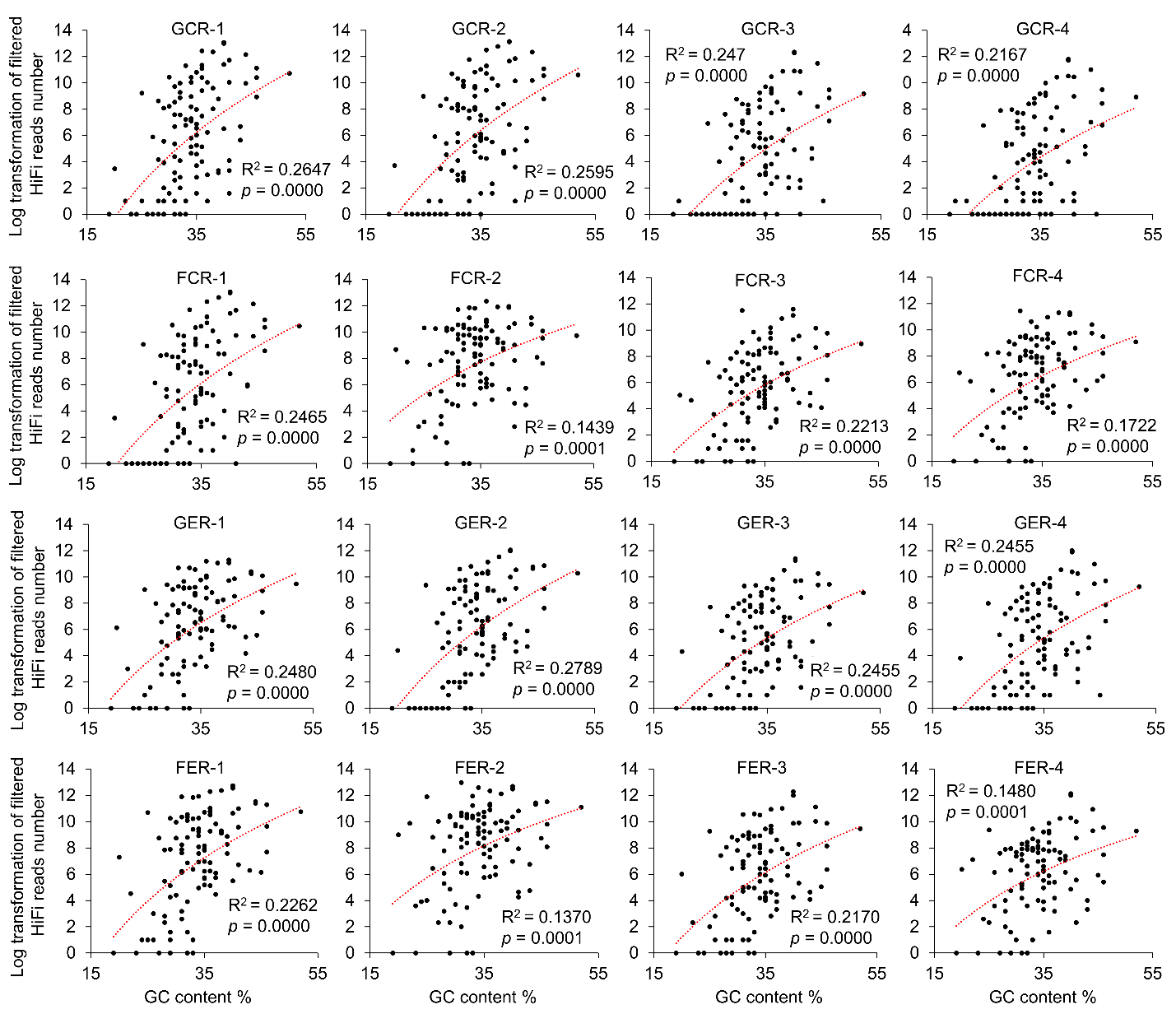

**Fig. S2.** Percentage of full-length reads containing all CG, CHG or CHH sites, and truncated reads not mapping 1 or more target sites, mapped in seven selected target regions for Ground control Col-0 Roots (GCR), Flight Col-0 Roots (FCR), Ground control *elp2-5* Roots (GER) and Flight *elp2-5* Roots (FER). For CHH sites, 1 site missing is allowed in the full-length reads.

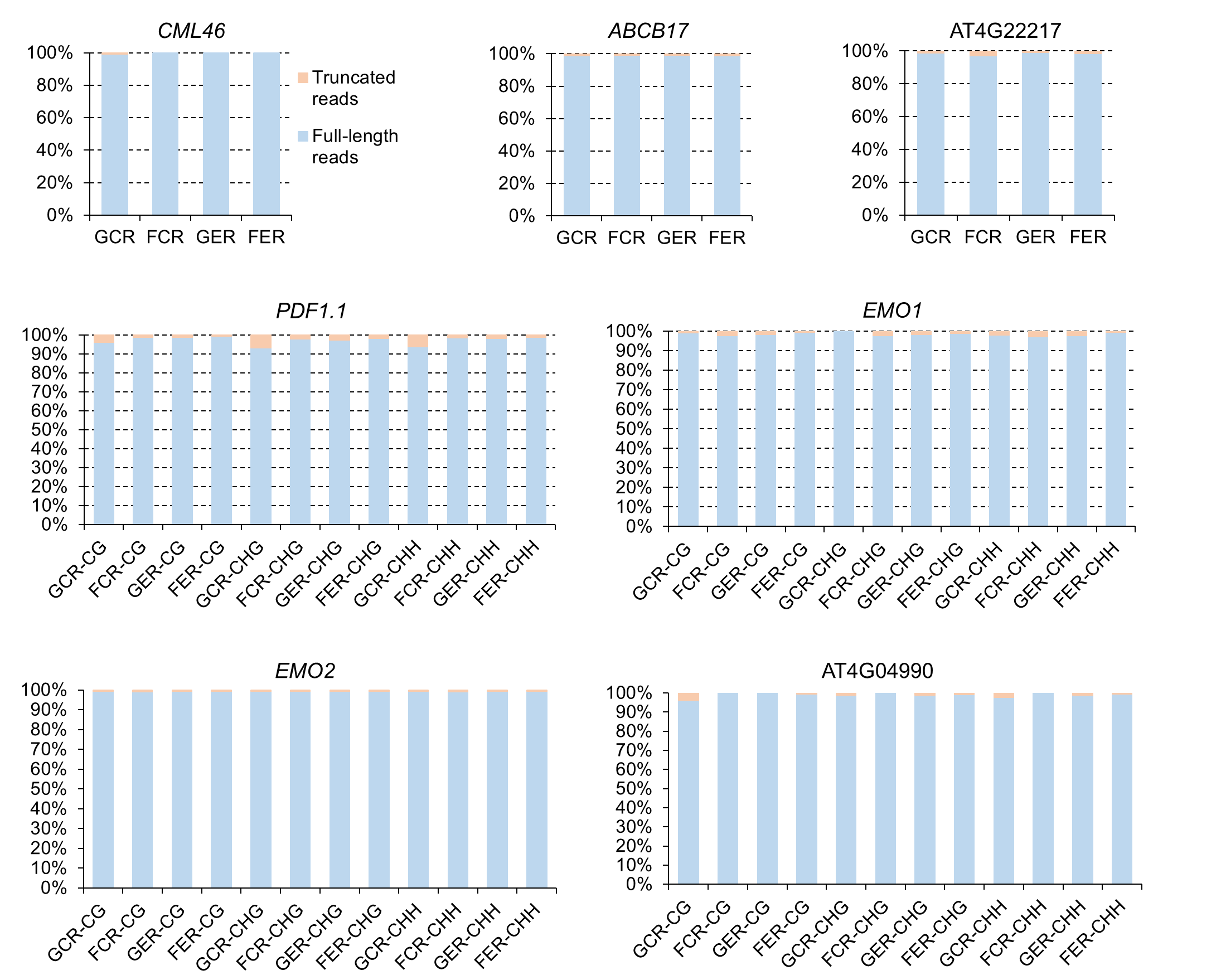

**Fig. S3.** FENGC results for *ABCB17*. (A) Genic region of *ABCB17*. The target region is indicated by brown rectangle (target ID: AT3G28380-1 in Table S1). The transcription start site is indicated by arrow. The captured fragment is in the coding strand of AT3G28380. (B) Expression levels of *ABCB17* that are presented in the same way of Fig. 2B. (C) Average CG methylation level within the target region. (D) Distribution of proportion of methylated CG sites per molecule in filtered HiFi reads containing all CG sites aligned to the target region. The significant differences of methylation proportion are shown (Kruskal-Wallis test; ***, *p* < 0.001). (E) Heatmaps showing the CG methylation on single molecules. The color legends are the same as Fig. 2E. The reads from the four biological replicates (S4 Table) were combined, then 1,000 reads were randomly selected to generate heatmaps for each sample. Approximately 0.8% reads in GCR, 3.8% reads in FCR, and 0.06% reads in FER containing at least 6 consecutive methylated CG sites are labeled by blue bricks. One-way ANOVA with post-hoc Tukey HSD test is performed for pairwise comparisons of percentage of reads with this footprint. FCR vs GCR, *p* < 0.05; FER vs FCR, *p* < 0.001.

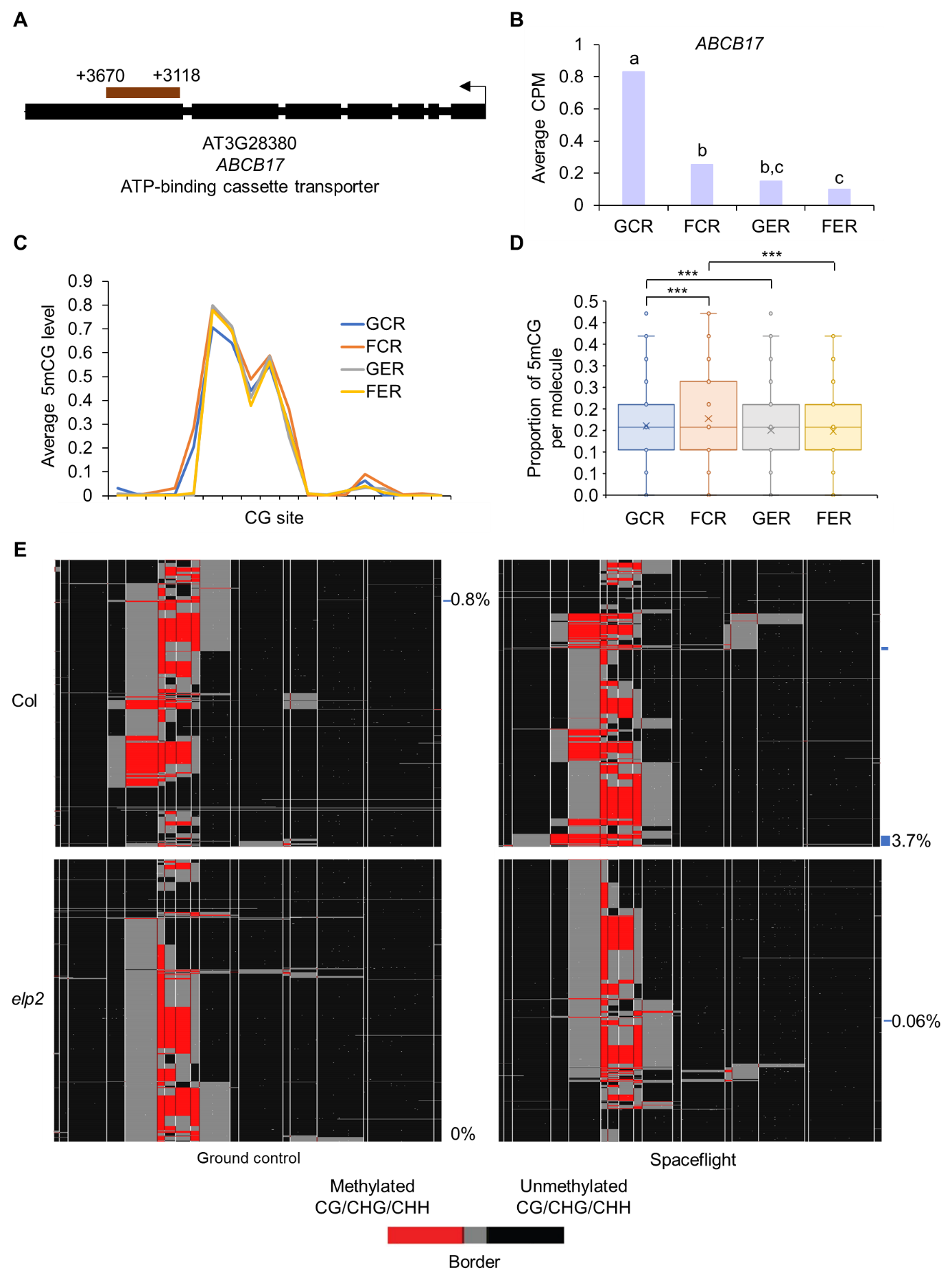

**Fig. S4.** FENGC results for AT4G22217. (a) Genic region of AT4G22217. The target region is indicated by brown rectangle (target ID: AT4G22217-1 in Table S1). A G-box and a MeJA-responsive element are indicated by yellow and green bricks, respectively. The transcription start site is indicated by arrow. (B) Expression levels of AT4G22217 that are presented the same way as Fig. 2B. (C) Average CG methylation level within the target region. The relative positions of *cis*-regulatory elements shown in (A) are indicated by yellow and green bricks. (D) Distribution of proportion of methylated CG sites per molecule in filtered full-length HiFi reads aligned to the target region. The significant differences of methylation proportion are shown (Kruskal-Wallis test; *, *p* < 0.05; ***, *p* < 0.001). (E) Heatmaps showing the CG methylation of single molecules. The color legends are the same as Fig. 2E. The reads from the four biological replicates (S4 Table) were combined to generate heatmaps for each sample. The relative positions of *cis*-regulatory elements shown in (A) are indicated by yellow and green bricks.

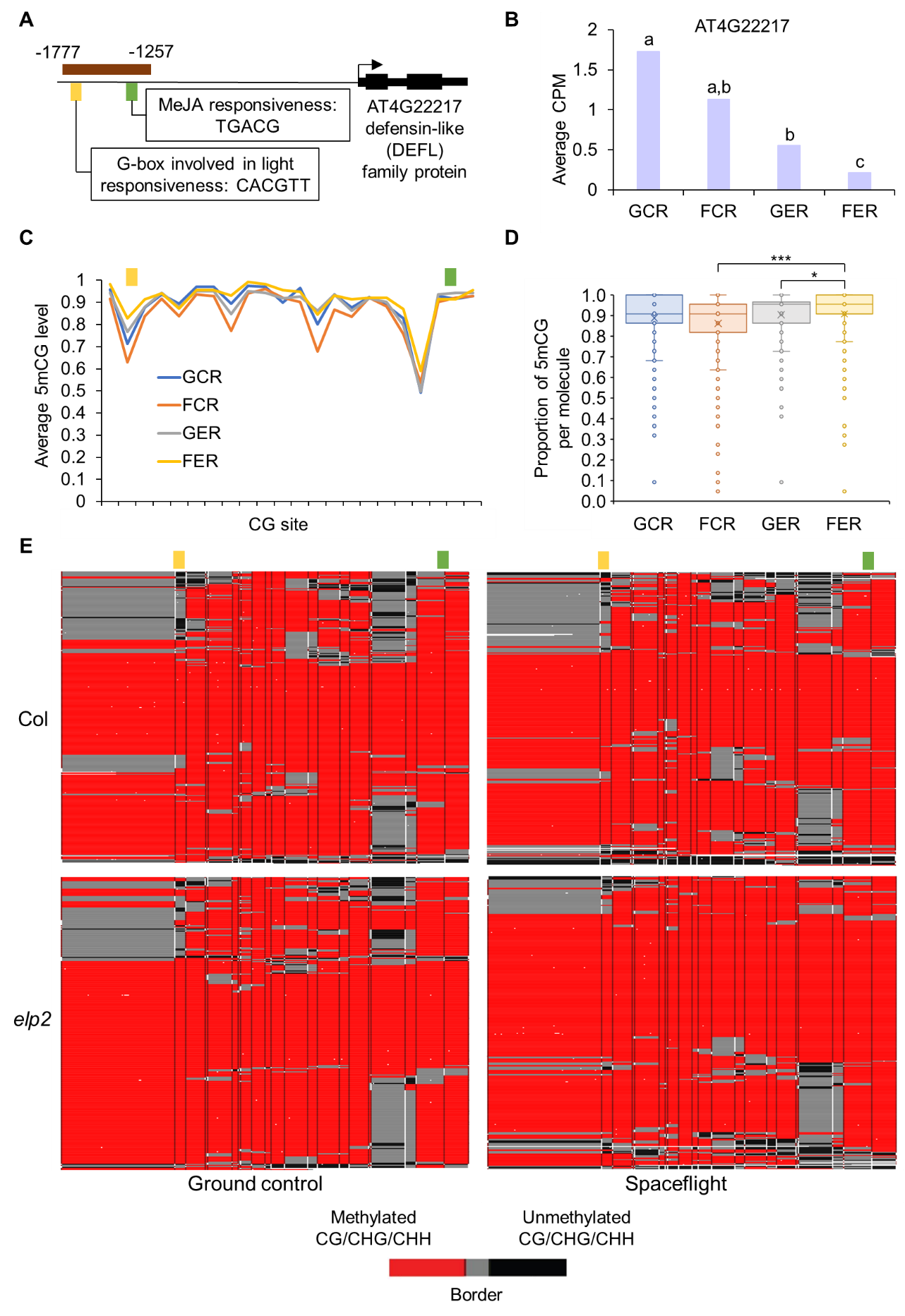

**Fig. S5.** Correlation between proportions of methylated cytosines in different sequence contexts within the same molecule in the captured region of *PDF1.1* (target ID: AT1G75830-1 in Table S1). R-squared value and *p* value for linear regression analysis are shown. Pairwise analysis revealed a trend of co-methylation across three types of cytosines in one cell population.

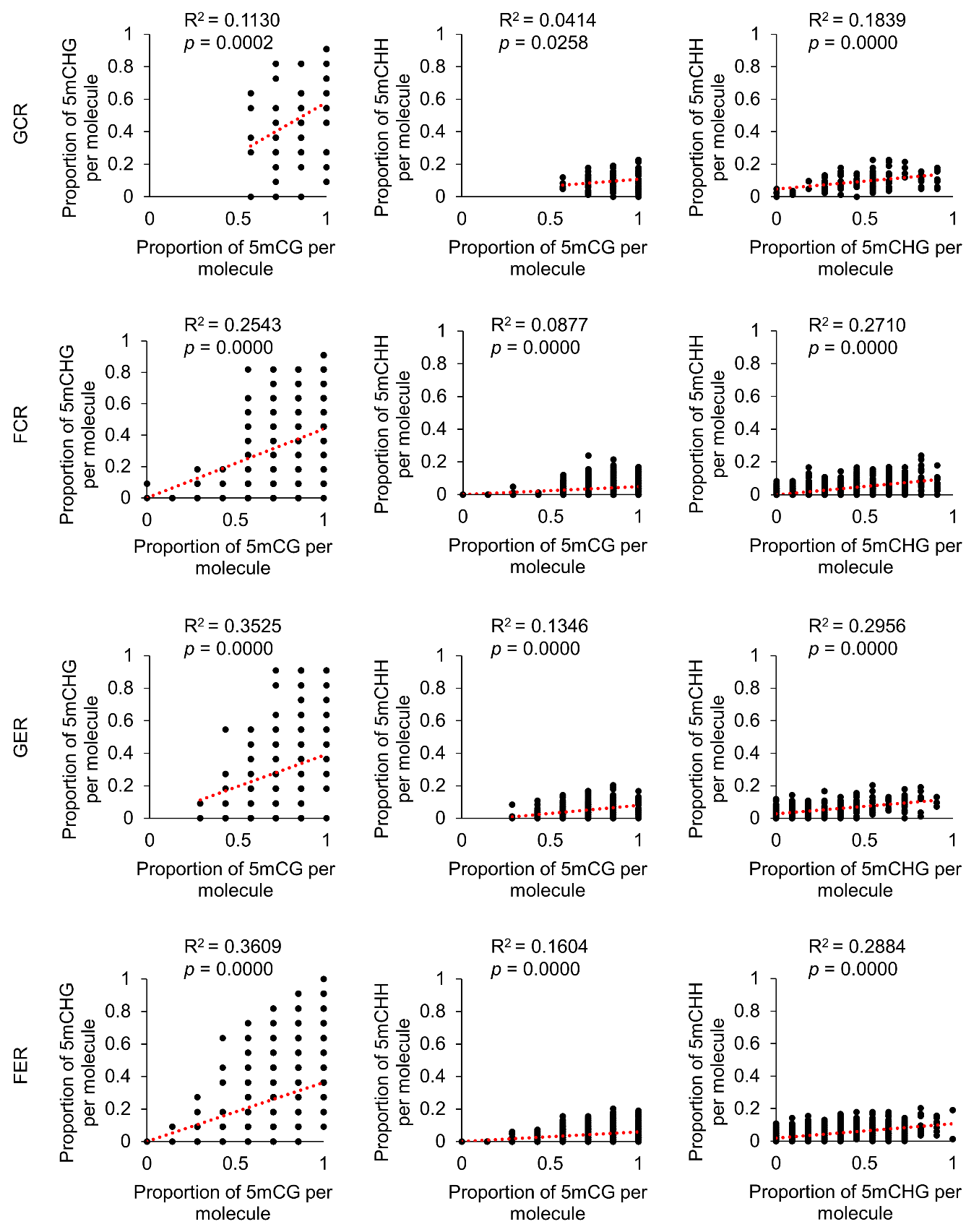

**Fig. S6.** Correlation between proportions of different types of methylated cytosines within the same molecule in the captured region of *EMO1* (AT1G27565). R-squared value and *p* value for linear regression analysis are shown.

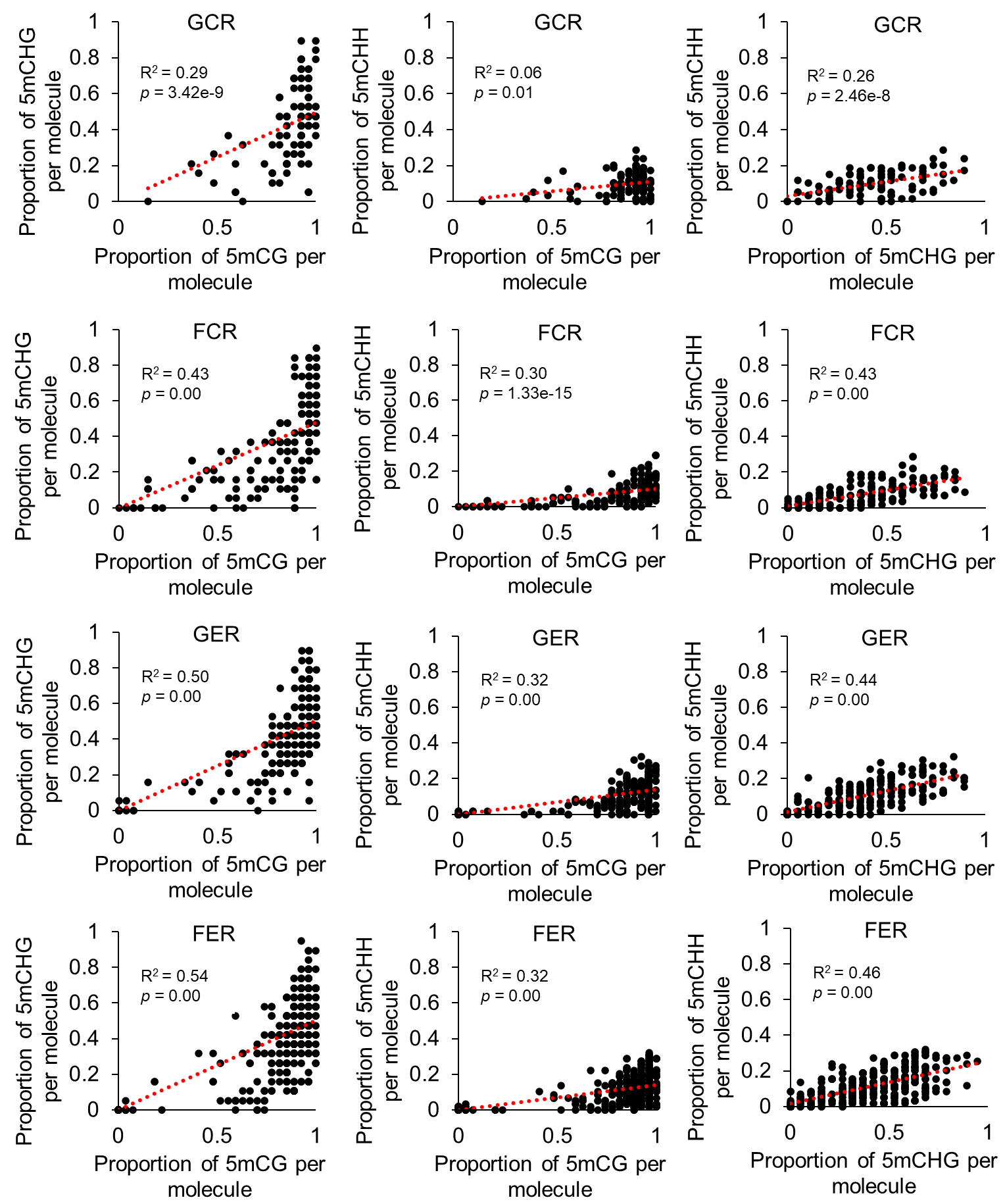

**Fig. S7.** Proportion of methylated CG, CHG and CHH sites within the same molecule in the captured region of *EMO1* (AT1G27565). There are 105, 181, 228 and 385 full-length reads covering all CG, CHG and CHH sites in this region, which are mapped for GCR, FCR, GER and FER samples, respectively. CG, CHG and CHH methylation proportions are indicated by three dots with different colors in the same column, which represents one read.

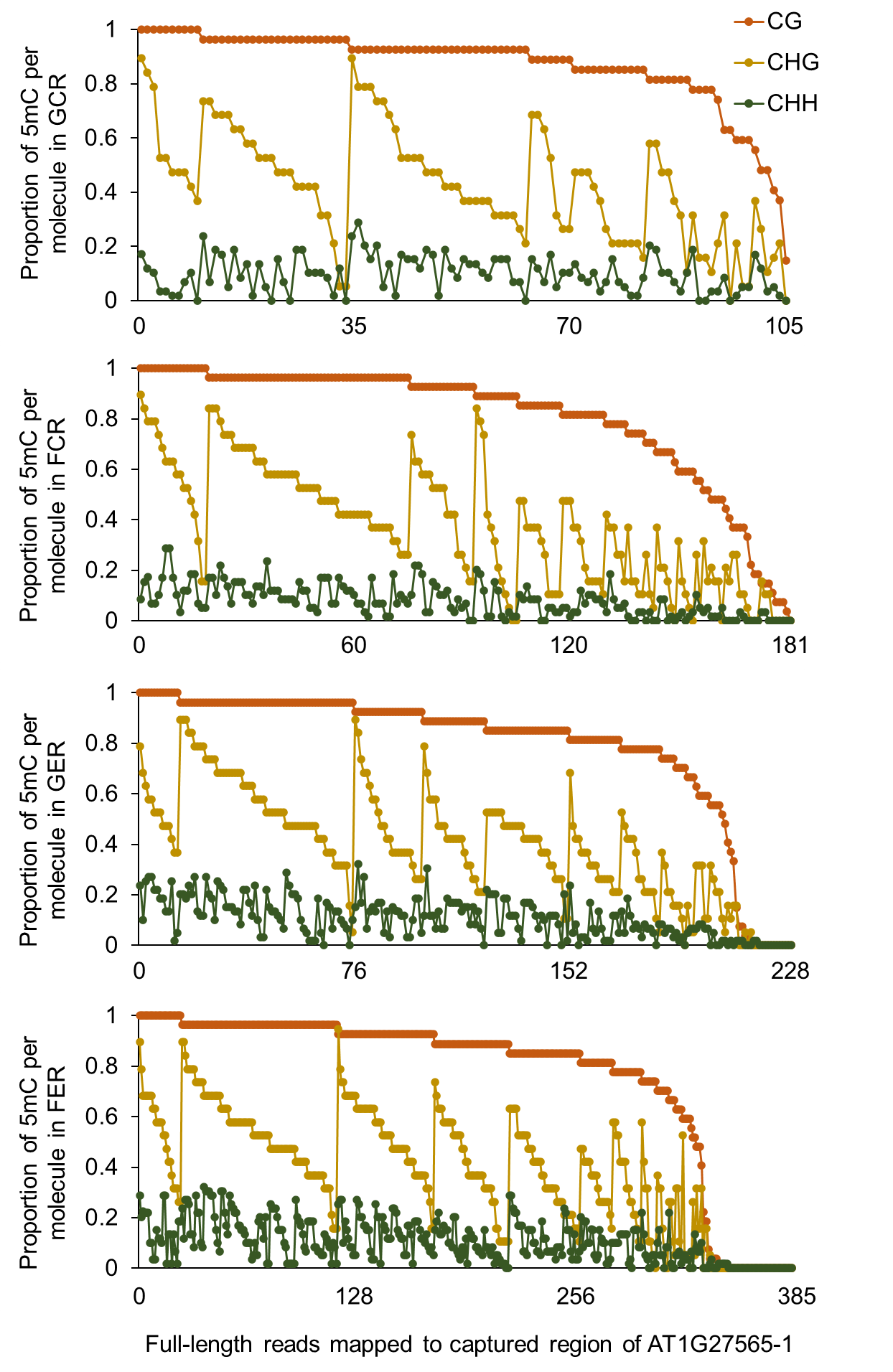

**Fig. S8.** Correlation between proportions of different types of methylated cytosines within the same molecule in the captured region of *EMO2* (AT5G52710). R-squared value and *p* value for linear regression analysis are shown.

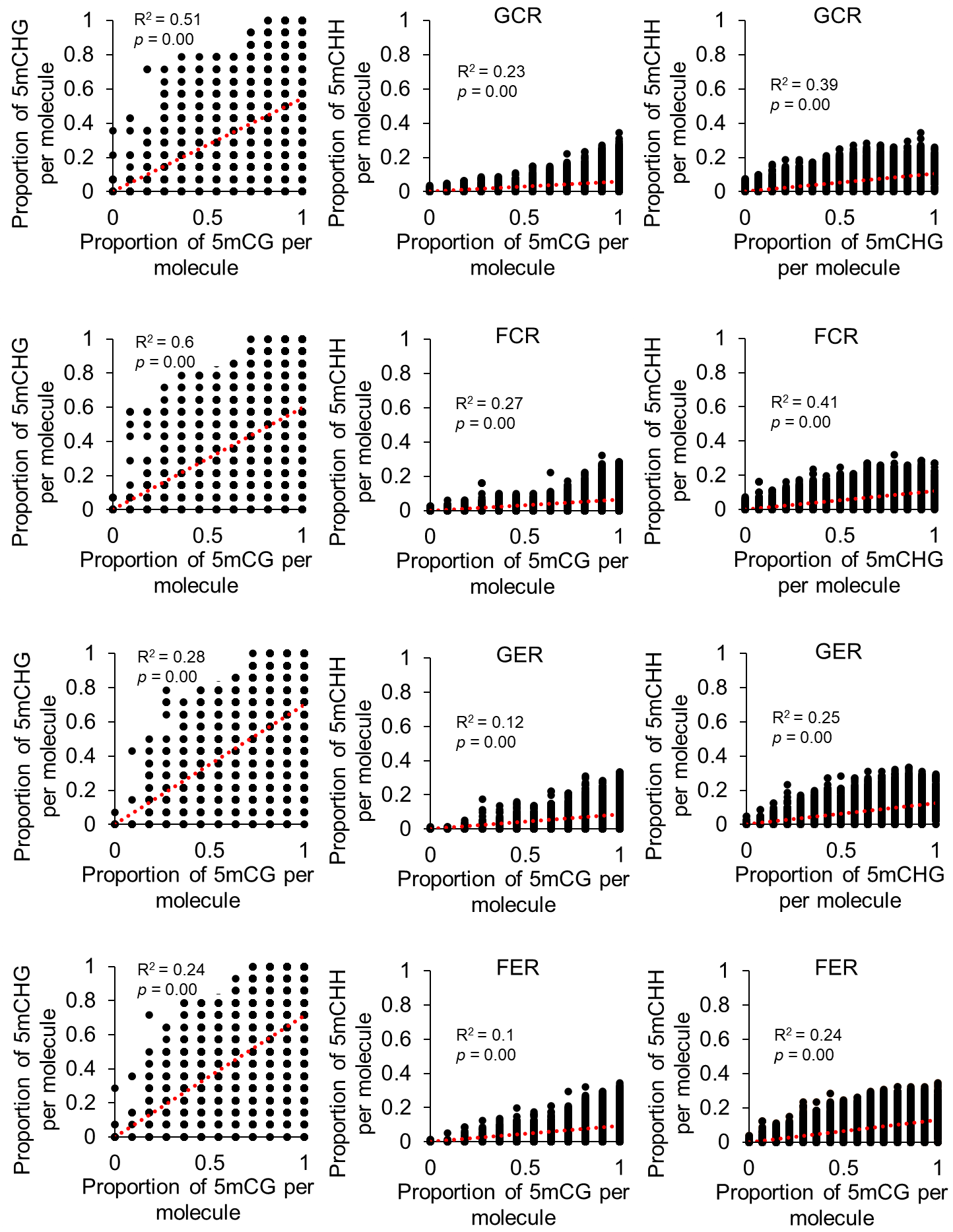

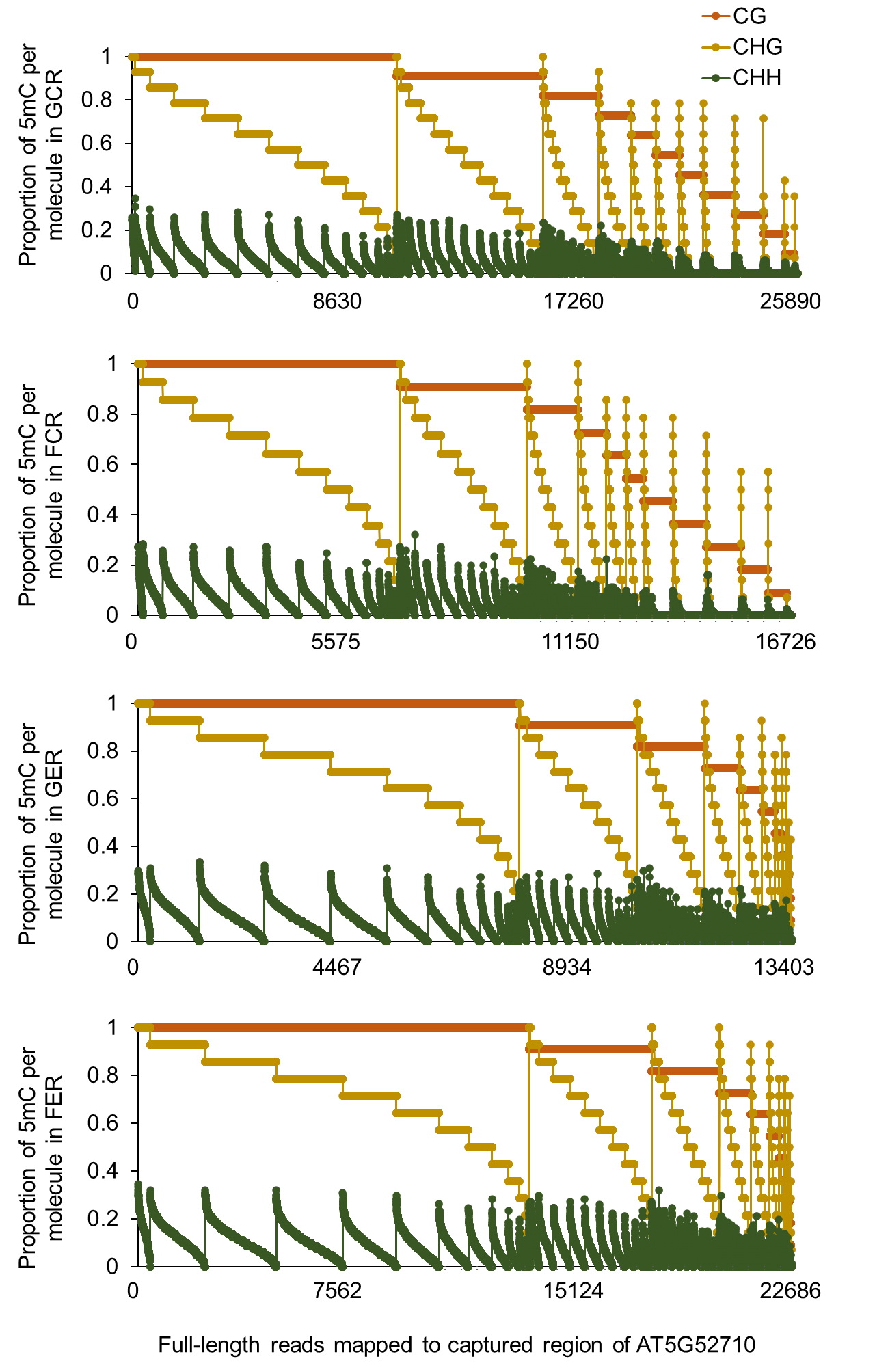
**Fig. S9.** Proportion of methylated CG, CHG and CHH sites within the same molecule in the captured region of *EMO2* (AT5G52710). There are 25890, 16726, 13403 and 22686 full-length reads covering all CG, CHG and CHH sites in this region, which are mapped for GCR, FCR, GER and FER samples, respectively. CG, CHG and CHH methylation proportions are indicated by three dots with different colors in the same column, which represents one read.

**Fig. S10.** FENGC results for AT4G04990. (A) Genic region of AT4G04990. The target region is indicated by brown rectangle (target ID: AT4G04990 in Table S1). A TC-rich repeat element is indicated by green brick. The transcription start site (TSS) is indicated by arrow. (B) Expression levels of AT4G04990 that are presented in the same way of Fig. 2B. (C) Average CG, CHG or CHH methylation levels in cytosines within the target region. The relative position of TC-rich repeats element shown in (A) is indicated by green brick. No significant difference of methylation levels is detected in the pair-wise comparison (One-way ANOVA with post-hoc Tukey HSD test). (D) Distribution of proportion of methylated CG, CHG or CHH sites per molecule in filtered full-length HiFi reads aligned to the target region. The significant differences of methylation proportion are shown (Kruskal-Wallis test; *, *p* < 0.05; **, *p* < 0.01). (E) Heatmaps showing the cytosine methylation in single molecules. The reads clustering, heatmap panels and color legend are the same as that in Fig. 5E.

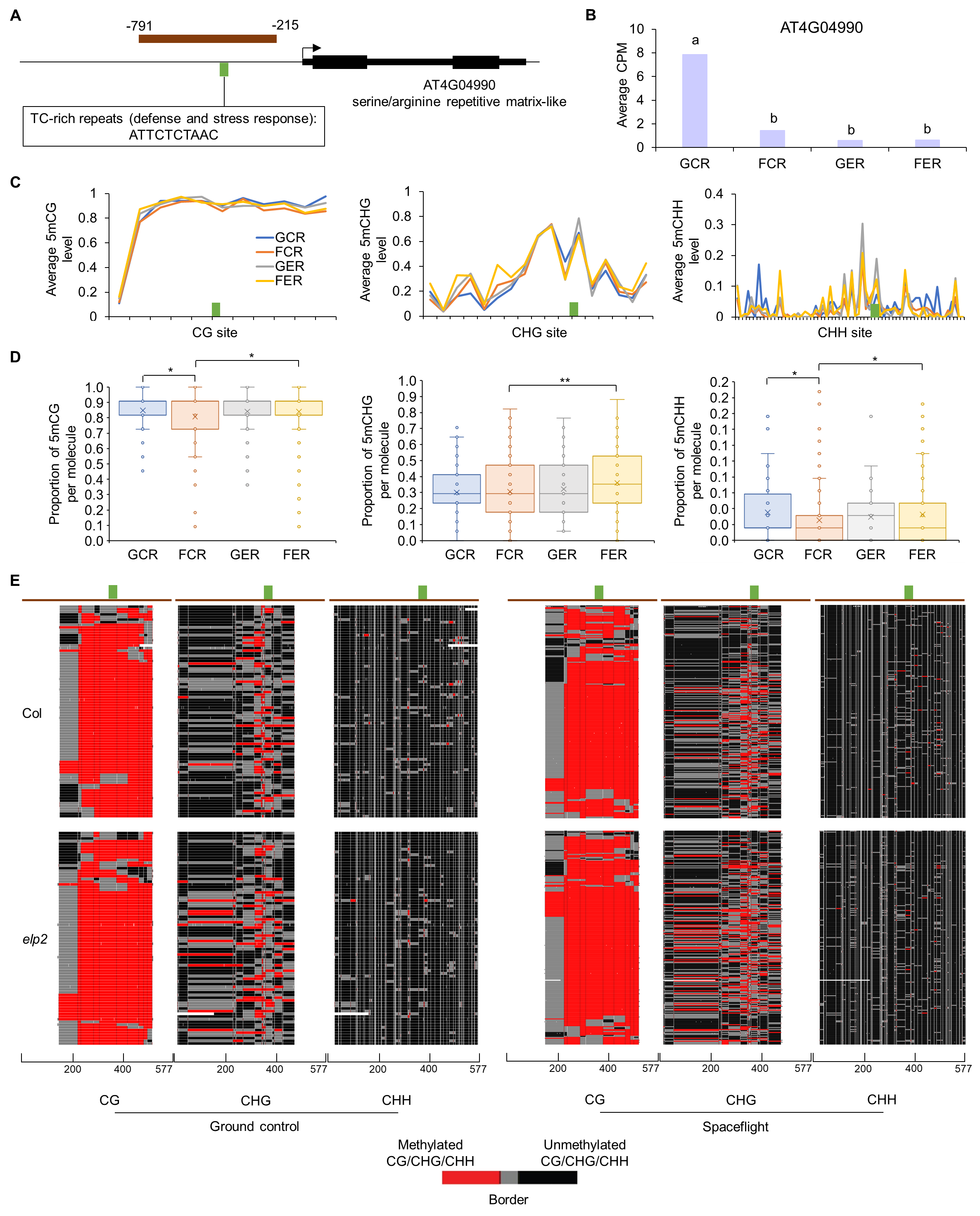

**Fig. S11.** Correlation between proportions of different types of methylated cytosines within the same molecule in the captured region of AT4G04990. R-squared value and *p* value for linear regression analysis are shown.

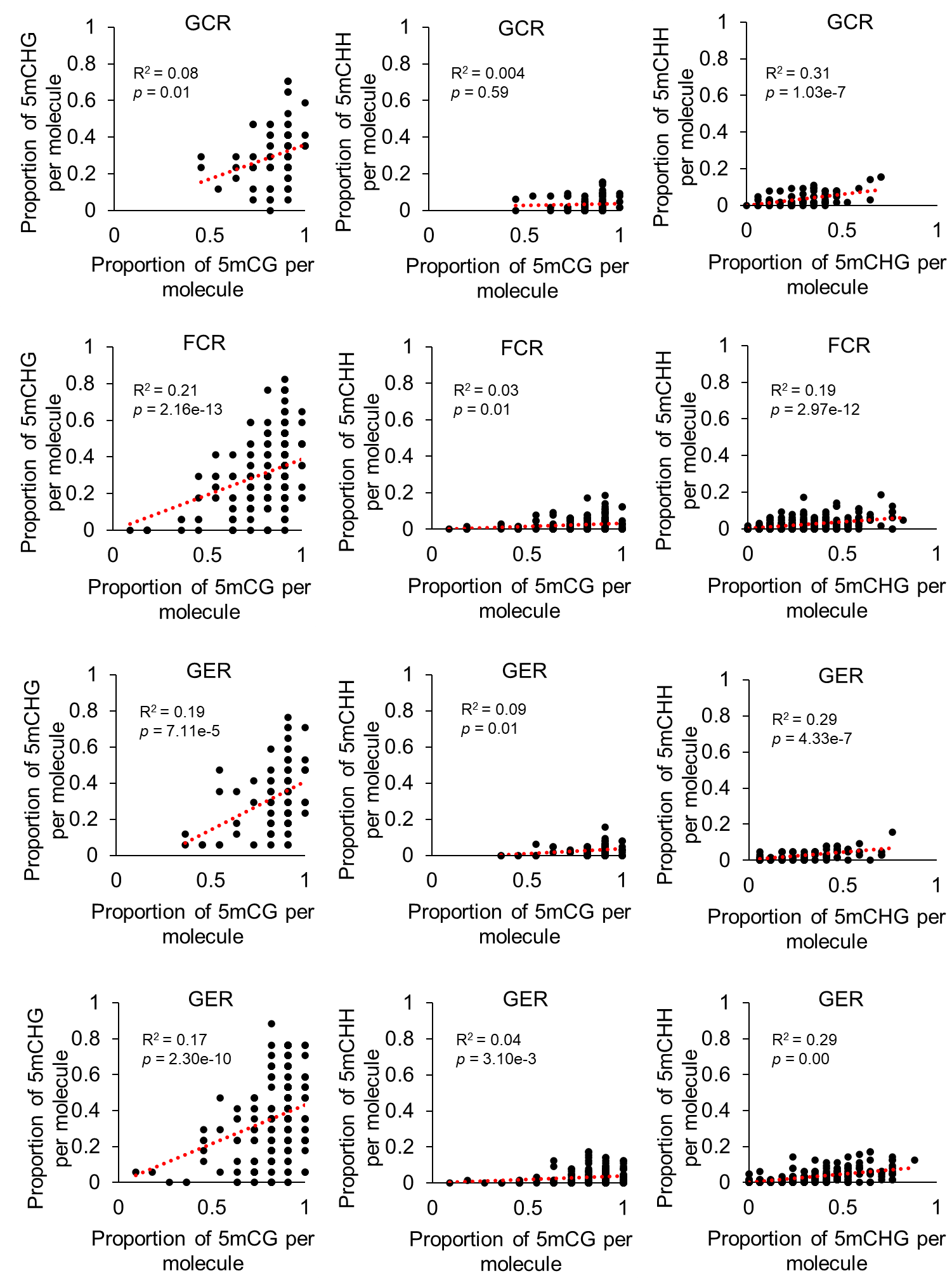

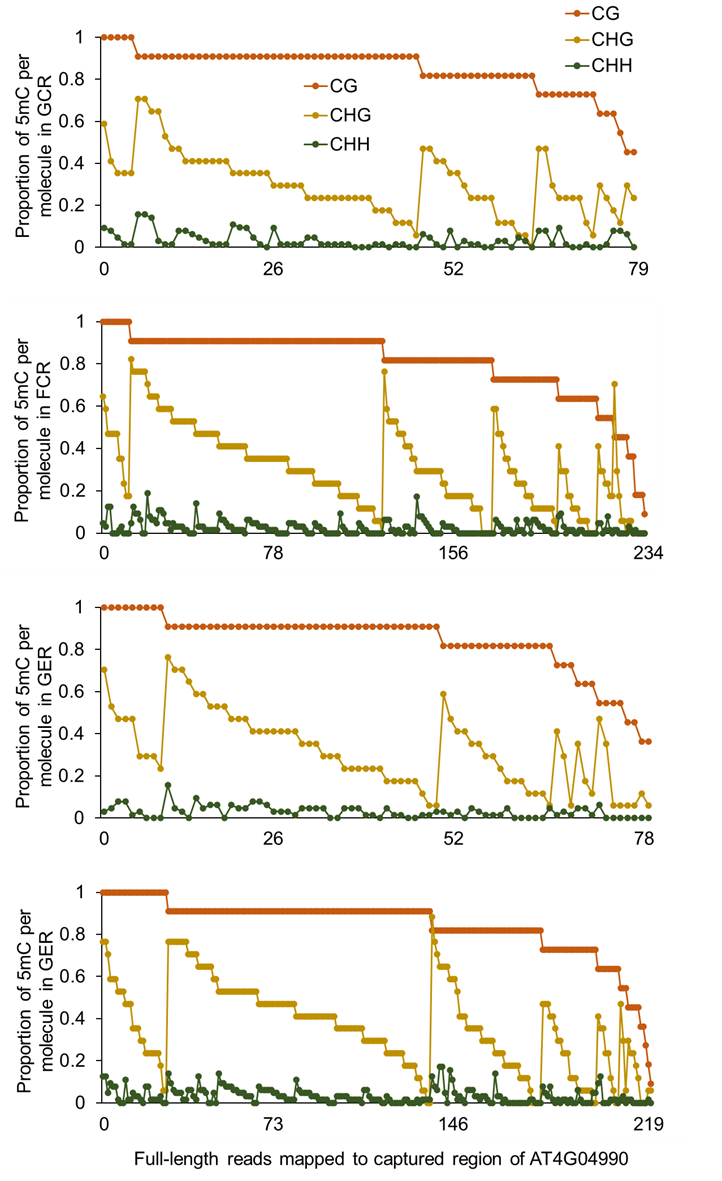
**Fig. S12.** Proportion of methylated CG, CHG and CHH sites within the same molecule in the captured region of AT4G04990. There are 79, 234, 78 and 219 full-length reads covering all CG, CHG and CHH sites in this region, which are mapped for GCR, FCR, GER and FER samples, respectively. CG, CHG and CHH methylation proportions are indicated by three dots with different colors in the same column, which represents one read.

**Fig. S13.** Relative transcription levels of *EMO1*, *EMO2* and AT4G04990 in their mutants, respectively. (A), (C), and (E), qPCR primers binding location in the genic structure. (B), (C) and (F), relative transcription levels. Two regions (R1 and R2) were used for *EMO2*. Data ± SE, n = 3.

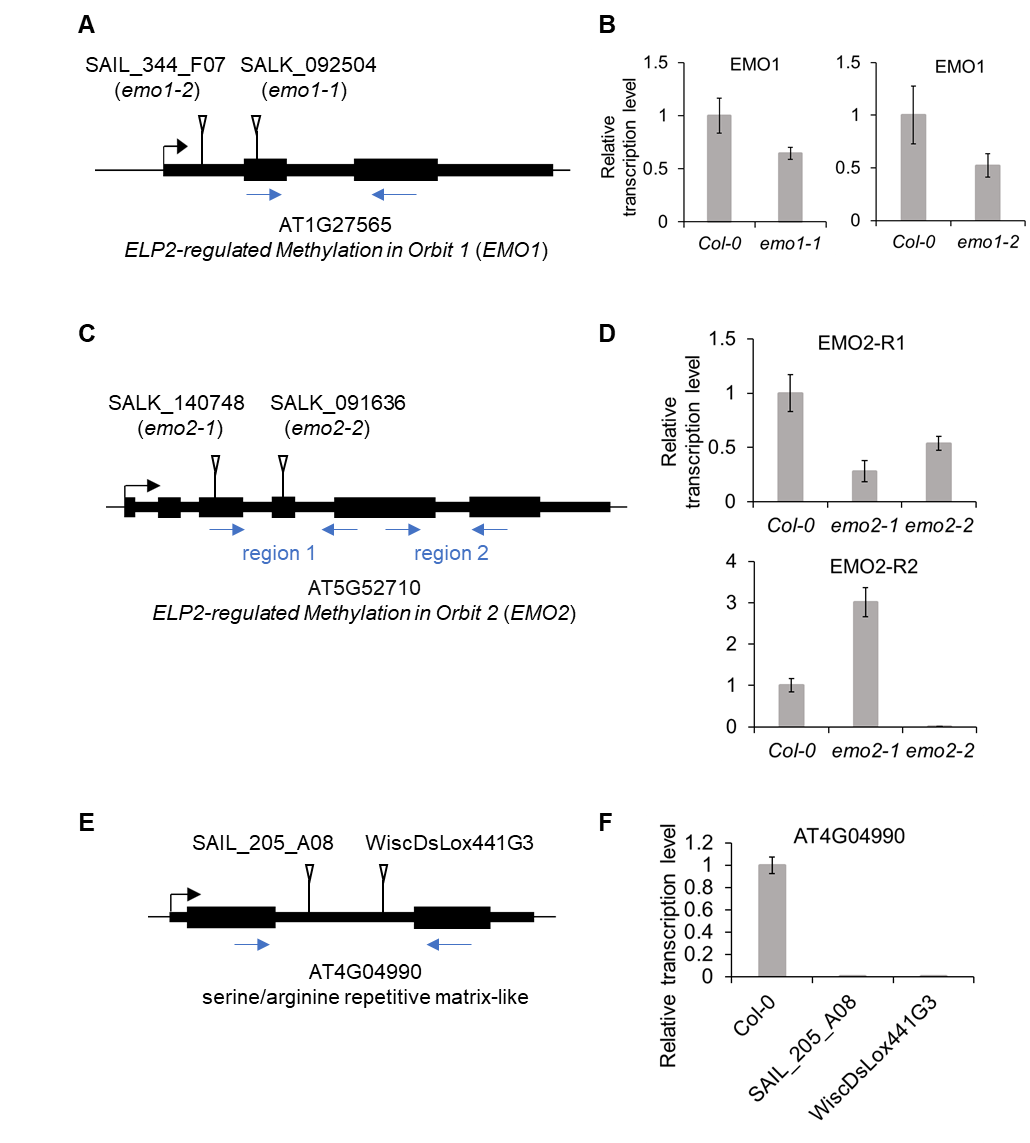

**Fig. S14.** Characterization of root growth in mutants of *EMO1*, *EMO2* or AT4G04990. (A) 7-day old seedlings of Col-0, SAIL_205_A08, and WiscDsLox441G3. Scale bars indicate 10 mm. (B) Primary root length of 4-day old and 7-day old seedlings. (C) Straightness (the ratio of straight distance from start to end of the root to root length). (D) Wave density (number of root bends per mm). (E) Skewing angle (angle from start to end of the root relative to the vertical). (F) HGI (Horizontal growth index, trigonometric relationship between the overall angle of growth and length of the root). For angles, positive and negative values indicate rightward and leftward growth of roots when viewed from behind the media, respectively. Data ± SD, n = 33 to 55 for each condition.

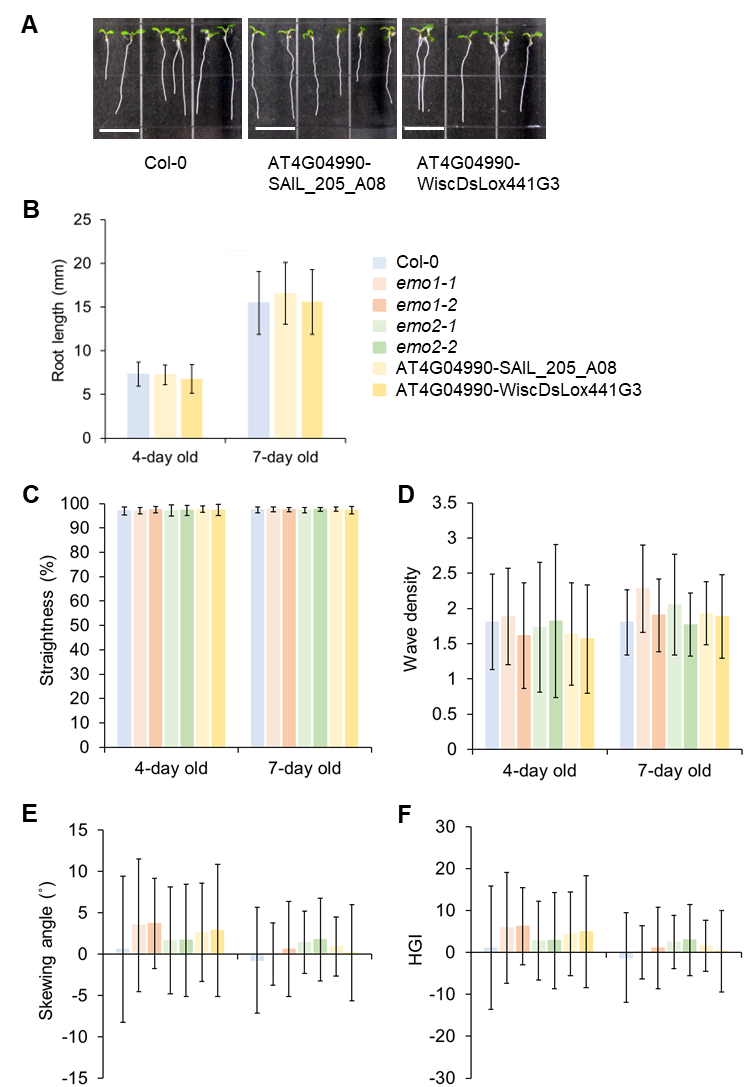

**Fig. S15.** Root gravitropism in mutants of *EMO1*, *EMO2* or AT4G04990. (A) The 4-day old seedlings growing in the vertical condition were transferred to a 2-dimentional clinostat and rotated at 4 rpm until 7-day old. (B) to (F) Primary root length, straightness, wave density, skewing angle, and HGI of 7-day old seedlings growing on clinostat. Student’s t test, *, *p* < 0.05; **, *p* < 0.01; ***, *p* < 0.001. Data ± SD, n = 8 to 23 for each condition. (G) The 4-day old seedlings growing in the vertical condition were reorientated 90˚ to the horizontal at time zero. Root tip angles from the horizontal were measured over time. Data ± SD, n = 11 to 19 for each condition.

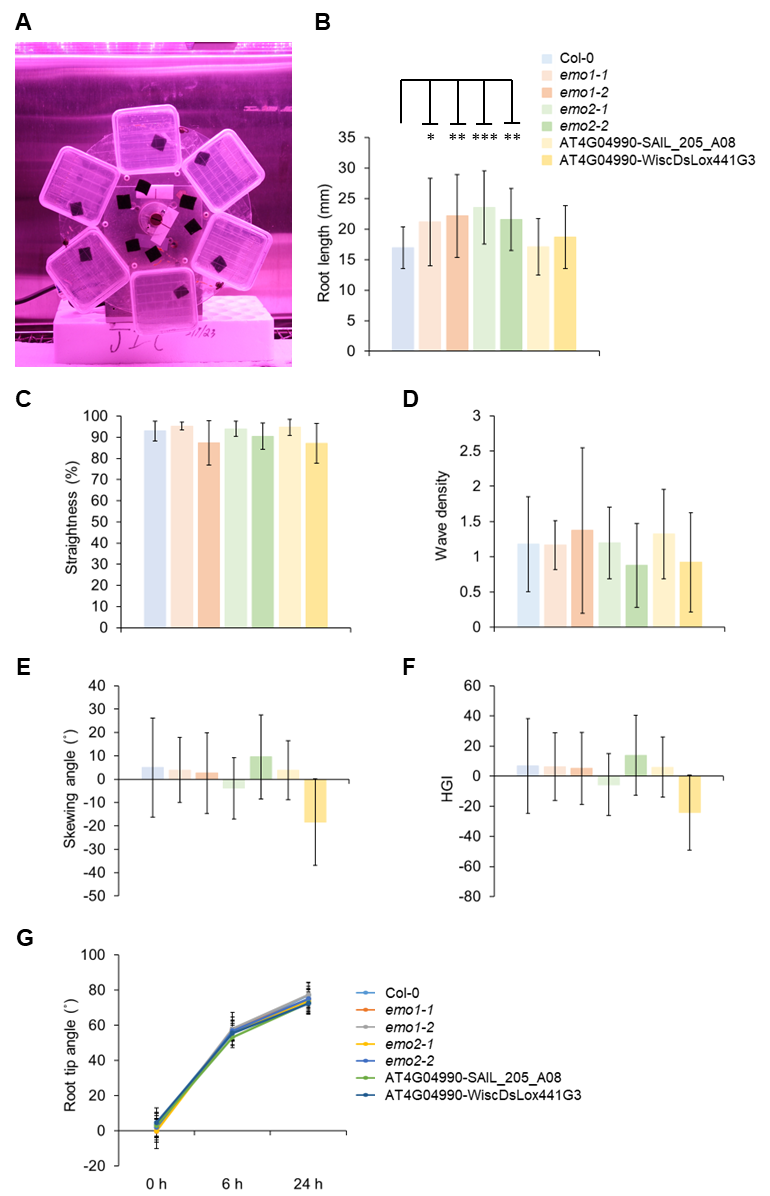
